# High Frequency Longitudinal RNAseq Reveals Temporally Varying Genes and Recovery Trajectories in Rats

**DOI:** 10.1101/2023.11.21.568082

**Authors:** Wei Chen, Yi Chai, Qi Jiang, Eva Y. Miao, Ashwin Gopinath, David Yu Zhang

## Abstract

When living organisms are exposed to potentially toxic chemicals, they respond via changes in gene expression. Traditional differentially expressed gene analysis based on before/after blood samples does not reveal the response’s temporal dynamics, and often produces false positives and negatives. Here, we performed longitudinal daily RNA sequencing on rats dosed with tetracycline, isoniazid, carbon tetrachloride, or valproate. We identified 4,302 temporally variable genes (TVGs) with statistically strong change in expression following dosing. Projecting TVG expressions into a 3-dimensional principal component (PC) space reveals consistent trajectories for recovery following dosing, and enables separation of healthy from recovering states with 91% to 99% area under the receiver operator curve. Finally, we observed that Fast Recovery vs. Slow Recovery rats exhibited distinct temporal expression patterns in the TVGs, suggesting that individual variations could be potentially captured via longitudinal RNAseq analysis.

---

Living organisms are dynamic systems; their RNA, protein, and metabolite concentrations change in response to changes in health and their environment. Of these, RNA expression is currently most affordable to comprehensively profile, due to advancements in high throughput sequencing (NGS) technology [1-3] over the past decade. By comparing the RNA expression profiles of different populations of individuals, researchers have identified unique RNA signatures for a variety of diseases and as predictive and prognostic biomarkers for therapy response [4-8]. This type of cross-sectional analysis allows identification of differentially expressed genes (DEGs) across two groups, but fails to capture the temporal dynamics of genes affected by perturbations to the individual, and also risks significant false discoveries [9] due to the noise intrinsic in the RNA sequencing (RNAseq) sample preparation and data collection process [10-11].

We hypothesized that high frequency longitudinal RNAseq analysis of blood samples would provide significantly deeper understanding of gene expression temporal dynamics in response to drug perturbations, and would also significantly improve the discovery accuracy of affected genes. In this study, we performed RNAseq on 829 blood samples collected from 84 Sprague-Dawley lab rats (*rattus norvegicus)*. Other than control animals, each rat was each dosed with one of 4 different small molecules (tetracycline, isoniazid, valproate, and carbon tetrachloride) with known liver toxicity. We developed custom bioinformatics analysis tools to identify temporally varying genes (TVGs).

## Experimental Design Summary

After acclimating the rats to their new environments for 3 days, we began daily blood draws, with Day 1 being defined as the first day of blood draws (Fig. 1a). Each blood draw was 200µL; this blood volume was selected as a balance between (1) obtaining sufficient biospecimen for analysis and banking and (2) not inducing undue harm to the rats from the bleeding process. The 200µL blood volume was decided considering of the potential accumulated effects of daily blood draws, and followed IACUC approved protocols. To ensure reproducibility of results and minimization of the effects of potential confounding factors, all blood draws were performed at approximately 10:00am local time each day. The relevant rats dosed with the respective drug intravenously on their Day 3 or Day 8, about 30 minutes prior to the next blood draw. Consequently, the first blood sample after dosing likely do not reflect the full effects of each molecule, based on the pharmacokinetics properties of each molecule. Following the end of each experiment, rat liver necropsy samples were analyzed via standard histopathology.

**Figure 1.**
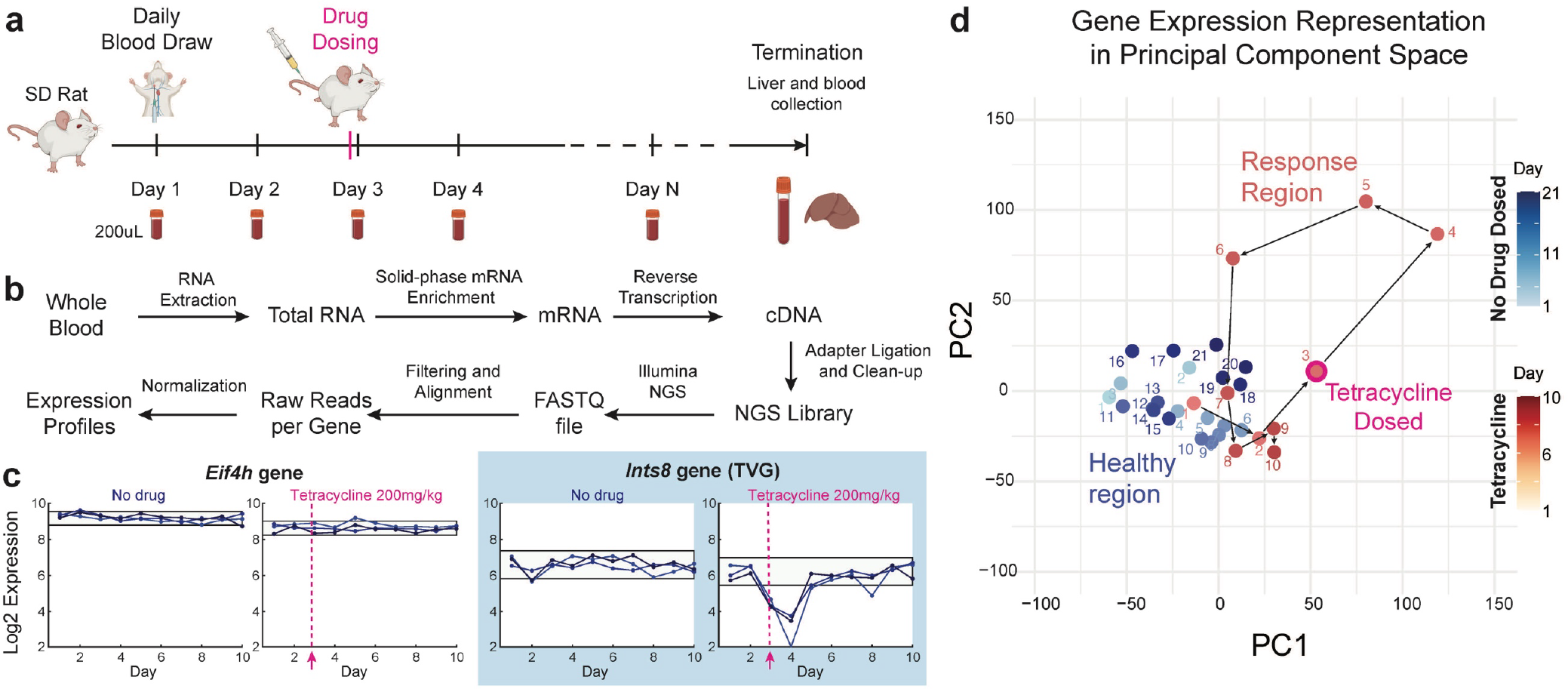
Overview of longitudinal rat RNAseq studies. (a) Animal experiment design. Sprague-Dawley rats are acclimated for 3 days at the contract research organization animal facilities before beginning daily blood draws (200 µL). Blood draws were performed at a consistent time of the day (around 10am) to minimize the impact of Circadian rhythm on gene expression. At least 2 daily blood draws were performed before dosing the animals with any drug molecules. (b) Workflow for mRNA sequencing and expression analysis. (c) Example longitudinal gene expression profiles for rats dosed with tetracycline just prior to Day 3 blood draw. (d) Trajectory of rat RNAseq results for control rats (blue) and in response to tetracycline dosing (red) in principal component space.

RNA was extracted from whole blood within 6 hours of collection. Subsequently, we used bead-based solid phase enrichment of mRNA from the total RNA sample using poly-T probes to remove non-coding RNA and ribosomal RNA. The enriched mRNA sample was then reverse transcribed using random hexamer primers, and the cDNA was sequenced using Illumina NGS at a depth of between 10M and 20M reads per sample. After filtering and alignment, we obtained the raw RNA expression profile, which was subsequently normalized based on the number of reads in each NGS library. See Supplementary Methods 1 and 2 for additional details on experimental and bioinformatics pre-processing methods.

As a preview of the data we collected and analyzed, Fig. 1c shows the temporal expression dynamics for two example genes, *Eif4h* and *Ints8*. Both genes showed temporally stable expression for the control rats, but the *Ints8* gene shows a 4-fold drop in expression on Day 4 for all 3 rats before returning on pre-dose levels on Day 5. Fig. 1d shows a projection of the tetracycline-dosed rats’ RNA expression on each data in a reduced 2-dimensional principal component parameter space. The trajectory of the rats through this parameter space shows that it took the rats roughly 4 days to fully recover from the dose, suggesting that the *Ints8* gene is an “early responder” gene. The dynamics of the rats’ recovery back to full health follows similar trajectories for all the molecules we tested, though we also noticed important individual variabilities in the magnitude of TVG RNA perturbation and the speed at which the rats recovered.

## Results

### Baseline and Bleeding Effects Characterization

Before conducting drug dosing experiments, we first performed RNAseq on daily blood draw for 21 days on N=3 male and N=3 female rats, in order to understand technical reproducibility, baseline gene expression variability, and the effects of daily blood draws on rats. This allows us to subsequently identify more confidently the genes whose temporal variations are due to the specific drugs being dosed.

Simple comparison of expression levels between pairs of rats on the same day confirmed the technical reproducibility of our results (Supp. Figs. S1 and S2). We next compared the RNA expression profiles of the 6 rats on Day 2 vs. Day 1. Because Day 1 was the first day that we drew blood, no bleeding or clotting related gene pathways are expected have been activated. Consequently, Day 1 data act as true baseline control RNA profiles. We used the standard DESeq2 analysis tool [12] to identify potential differentially expressed genes (DEGs) in Day 2 vs. Day 1 (Fig. 2a). The standard p=0.05 significance cutoff for calling DEGs resulted in many false positives (Supp. Fig. S3) and was insufficient to identify true DEGs.

**Figure 2.**
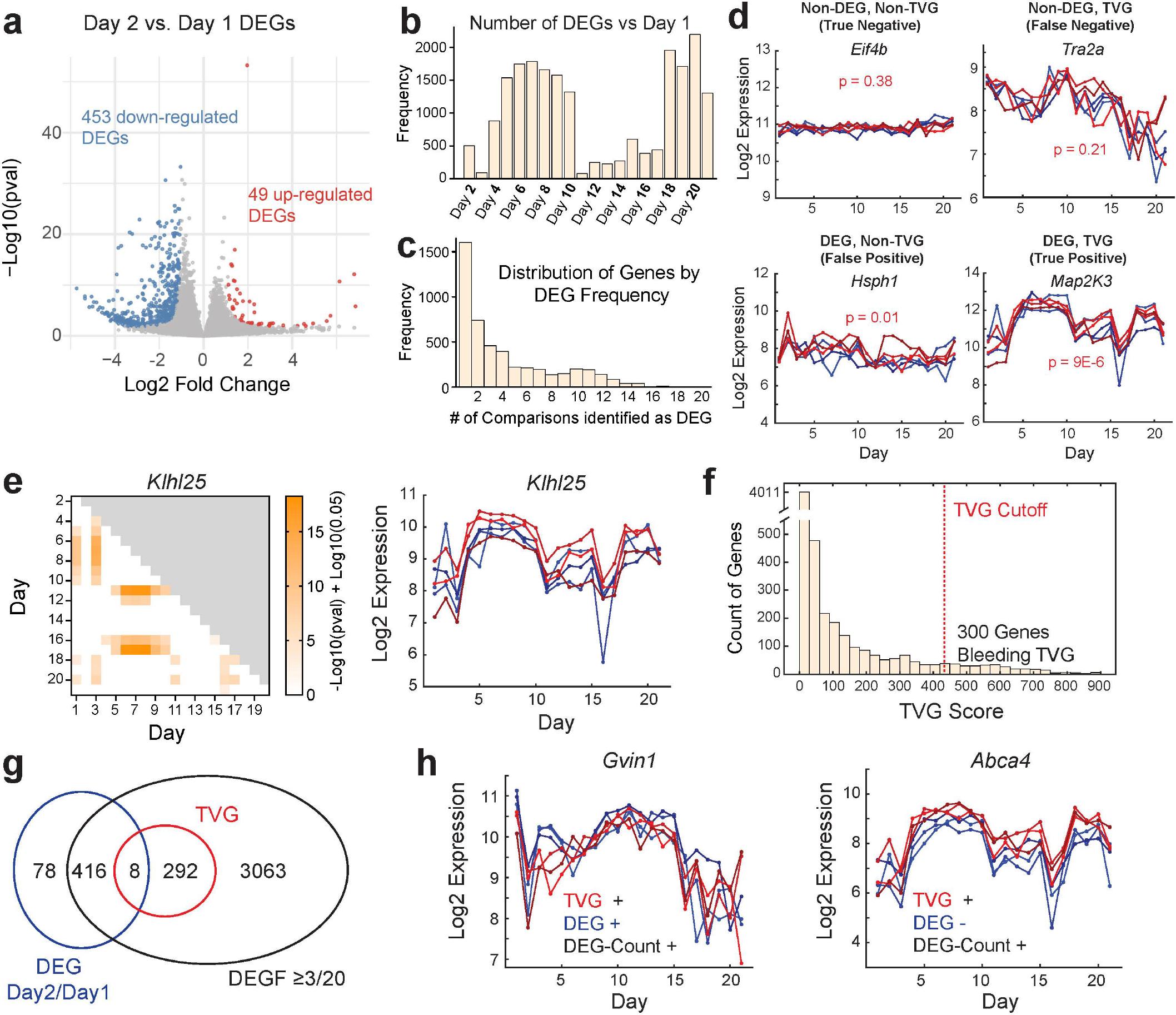
Control studies and impact of daily blood draws. (a) Analysis of differentially expressed genes (DEGs) using DESeq2, comparing Day 2 vs. Day 1 for N=6 rats. A total of 502 DEGs were identified, based on our filters (See Supp. Excel 1). (b) Analysis of longitudinal RNAseq data, using Day 1 expression profile as a reference. We identified significantly more DEGs in Days 5-9 and Days 18-21 than Days 11-17, suggesting a potential biological effect with 2-week periodicity. (c) Distribution of number of genes differentially expressed on multiple days vs. Day 1. We identified 3779 genes with DEG frequency (DEGF) ≥ 3, based on comparisons to Day 1. (d) False positive and false negative generated by DEG analysis considering only Day 2 and Day 1. The *Tra2a* gene has statistically significant expression decrease over time, but is not flagged as a DEG due to the large variance of gene expression on Day 2. In contrast, the *Hsph1* gene shows high expression with low variance on Day 2, but overall appears to be relatively stable in expression, suggesting a potential false positive from DEG analysis considering only 2 timepoints. (e) Our framework for identifying Temporally Varying Genes (TVGs) based on p-values of the pairwise DEG analyses. The *Klhl25* gene shown here is the gene with 300th highest TVG score, and the most marginal gene included as a TVG. (f) Distribution of TVG scores. The top 300 genes were selected as bleeding/baseline TVGs for downstream analysis. (g) Venn diagram comparing TVGs vs. genes identified as DEGs based on pairwise comparison. (h) Example TVGs gene expression over time.

To improve the accuracy of DEG calling, we developed and used a metric comprising a non-linear combination of adjusted p-value and log2 fold change, with a single fitting parameter k (Supp. Fig. S4).

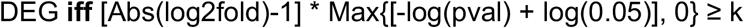

Setting k=1 removed the vast majority of false positive findings from DESeq2 analysis (gray dots in Fig. 2a). Using this approach, we identified 49 up-regulated and 453 down-regulated DEGs in Day 2 (Supp. Excel E1). These 502 genes correspond to roughly 2.1% of all 23,987 rat genes, and roughly 5.6% of the 9,044 rat genes with significant non-zero expression in our studies.

The 21-day longitudinal blood RNAseq data that we collected allows more sophisticated analysis and identification of TVGs. We first attempted to identify TVGs through DEGs identified via pairwise comparison of each Day’s RNAseq profiles against the Day 1 data (Fig. 2b). Unexpectedly, we found that the number of DEGs varied by more than 10-fold, depending on the Day that is being analyzed. Furthermore, the number of DEGs appeared to follow a smooth pattern with a 2-week period, rather than being independently distributed. Visual analysis of temporal gene expression confirmed the periodic behavior of gene expression for a subset of genes (*Tra2a* and *Map2k3* in Fig. 2d). To the best of our knowledge, this is the first experimental report of such time-dependent gene expression phenomenon at the scales that we have observed (affecting more than 1000 genes). We are unsure of the exact nature of these genes, as female rats do not menstruate in the way humans do and these gene temporal dynamics were also present in male rats.

We next considered the number of days that each gene’s out-day expression significantly varied from Day 1 expression (Fig. 2c). We decided to test the definition of TVGs being those genes that are called as DEGs with frequency grater than ≥3 out of 20 (DEGF ≥ 3), based on the 20 pairwise comparisons vs. Day 1. Comparing the genes with DEGF ≥ 3 against the DEGs called from Day 2/Day 1 showed that the latter generated many false positives and false negatives (Fig. 2d). False positives could be partially due to measurement noise (e.g. *Hsph1*), and false negatives could be due to delayed activation or periodic expression genes not significantly different in expression across consecutive days. These findings highlight the importance and contribution of high frequency longitudinal RNAseq collection.

Visual spot check inspection of the temporal expression profiles of the genes with DEGF ≥ 3 suggested, however, that this approach likewise generated significant false positives, primarily because Day 1 data was overweighed. After considering and evaluating multiple approaches for defining TVGs, we decided on a TVG score based on the p-values of DEG analysis across all pairwise comparisons across different Days.

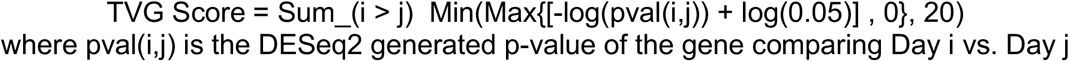

Fig. 2e shows the contributing p-value components of the *Klhl25* gene, and its temporal gene expression profile (see also Supp. Fig. S5), and Fig. 2f shows the distribution of TVG scores for all expressed rat genes. Based on visual inspection of the temporal gene expression profile of the 6 rats, we decided that the 300 genes with highest TVG scores reasonably qualified as true TVGs. All 300 of the TVGs identified in this way exhibited DEGF ≥3, but only 8 of them were identified as DEGs by comparing Day 2 vs. Day 1 (Fig. 2g). These observations suggest that a large majority of the TVGs identified fluctuate naturally or are delayed responses to bleeding, as opposed to acute bleeding responses. Fig. 2h shows the gene expression profiles of two TVGs. Of these, *Gvin1* is also identified as a Day 1 / Day 2 DEG and *Abca4* is not. See Supp. Figs. S6-S10 for additional examples of TVG expression profiles, and Supp Excel E2 for the full list of called TVGs.

### Liver Toxicity Responses

We dosed the rats with 4 different small molecules: tetracycline, isoniazid, valproate, and carbon tetrachloride. The first 3 are FDA approved drugs, and the last is a formerly used drug (Necatorina) that is now a standard molecule for testing liver damage. In our studies, we found that the highest number of TVGs in response to tetracycline dosing, and will primarily display tetracycline results in the main text and figures. See Supplementary Excel E3 for experimental design.

We dosed N=3 rats at each of 4 different concentrations of tetracycline (4, 15, 50, and 200 mg/ kg), and unsurprisingly the highest dose elicited the strongest gene expression response. Consequently, we used the 200mg/kg dataset to do TVG identification, following the same method as in Fig. 2ef. In this analysis, we excluded the 300 bleeding/baseline TVGs that we previously called, and identified 3,206 TVGs for associated with tetracycline response (Fig. 3a). Temporal expression profiles for the TVGs *Psmb4* and *Tpm4*, as well as non-TVG *Grk6*, are shown in Fig. 3b. *Tpm4* responds quickly to tetracycline, with the strongest gene down-regulation on Day 4. Psmb4, in contrast, shows the highest up-regulation on Day 5, suggesting potentially either delayed gene activation or slower deactivation.

**Figure 3.**
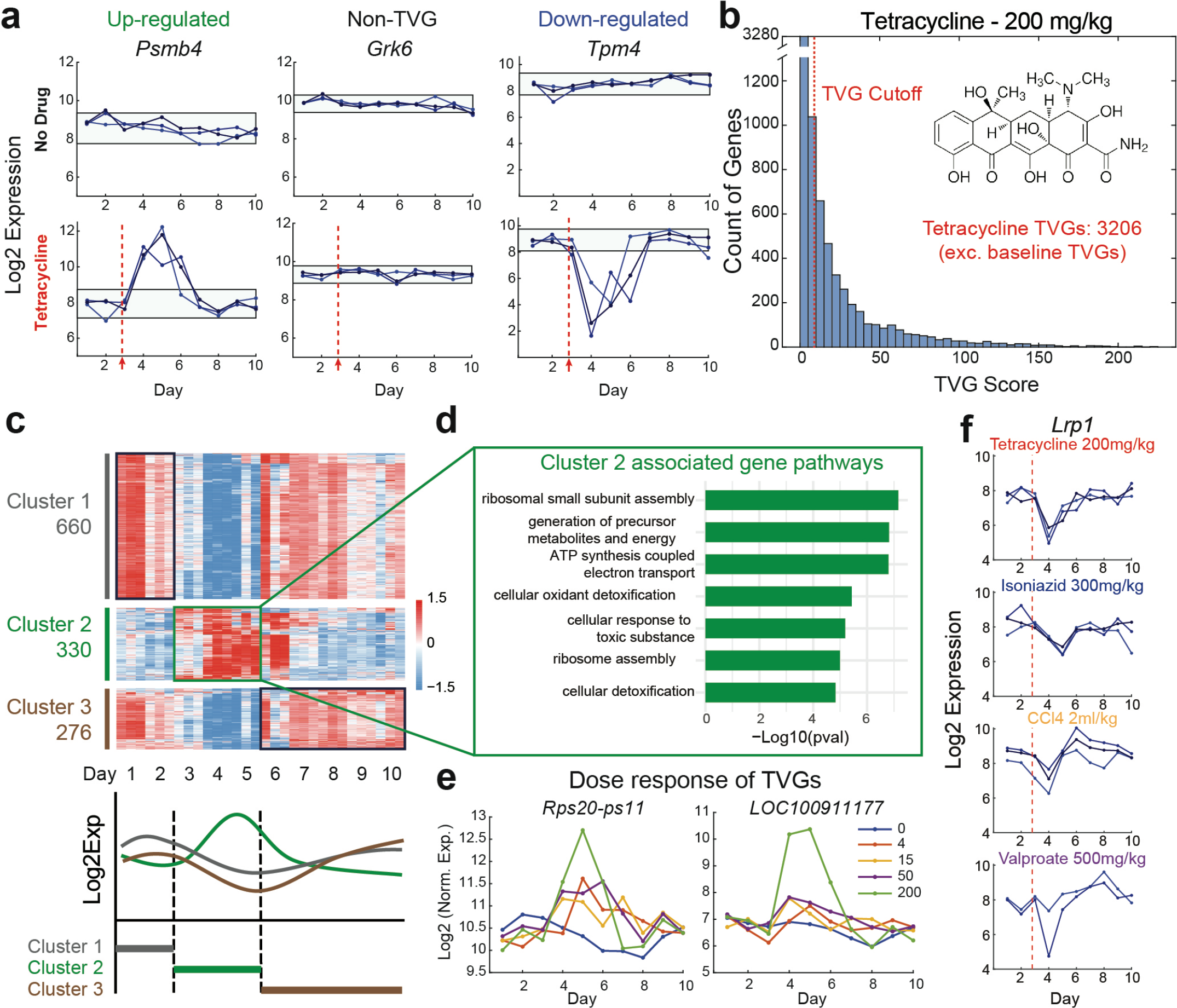
Rat response to tetracycline and other drugs. (a) Example up-regulated TVG, non-TVG, and down-regulated TVG for tetracycline. Horizontal lines show +/-2 standard deviations from mean log Expression. (b) Summary of TVG scores for 200mg/kg tetracycline. 3206 TVGs were identified. (c) Unsupervised clustering the tetracycline TVGs produces 3 clusters of genes with different temporal dynamics. Genes with 0 expression or less than 0.5 Log2 fold change from mean were excluded from analysis. (d) Pathways analysis of cluster 2 TVGs (up-regulated in Days 3-5) shows statistically significant association with cellular response to toxic substances. (e) TVGs identified from 200mg/kg tetracycline had different dose responses. *Rps20-ps11* was upregulated even at the lowest dose of 4mg/kg, while *LOC100911177* was upregulated only for the highest 200mg/kg dose. (f) We also performed TVG analysis on 3 other drugs (isoniazid, valproate, and CCl4). Shown here is *Lrp1*, a TVG for all 4 drug molecules.

To identify the function of the genes, we perform clustering to group the TVGs based on their temporal expression profiles, focusing on the genes with significant expression changes vs. the mean on Days 1-2, Days 3-5, or Days 6-10 (Fig. 3c). Of these three clusters of TVGs, Cluster 2 showed upregulation on Days 3-5, suggesting potential role in liver damage recovery. Analysis of the Cluster 2 TVGs using ClusterProfiler [13] showed several groups of associated gene pathways. See Supp. Fig. S11 for pathways associated with Clusters 1 and 3 for tetracycline, Supp. Fig. S12 for pathways associated with TVGs for isoniazid, valproate, and CCl4. To further confirm that the associated gene pathways are real response signals, rather than potential false positives from testing a large number of gene hypotheses, we examined the impact of tetracycline dose on the magnitude of the gene response (Fig. 3e). Consistent with the up-regulated TVGs contributing to drug response, we found that the up-regulated TVGs had weaker responses at lower doses. Furthermore, we noted that different TVGs were activated at different drug doses: e.g. *Rps20-ps11* was up-regulated on Day 5 even at the lowest dose of 4mg / kg, but *LOC100911177* was up-regulated only at the highest 200mg/kg dose. See Supp. Fig. S13 for statistics on the tetracycline dose activation threshold.

Analyzing the other 3 drug molecules, we identified 1,742 TVGs for isoniazid, 1,233 TVGs for carbon tetrachloride, and 1,068 TVGs for valproate. The union of these TVGs with the 3,206 tetracycline TVGs yielded a total of 4,302 TVGs. Of these, 186 genes were identified as TVGs for all 4 drugs, including the *Lrp1* gene (Fig. 3f). Given the large number of TVGs identified specific to different subsets of drugs, it is likely that these drugs activate multiple different pathways for recovery. See Supp. Fig. S14 for additional analysis of TVG overlap, and Supp. Figs. S15-S18 for summary and gene clustering analysis for the 4 drug molecules.

### Principal Component Space Trajectories

In addition to our bottom-up analyses thus far, we also aimed to see what macroscopic insights could be obtained from the longitudinal RNA expression data from the top down. We took the 4,302 TVGs (for any of the 4 molecules) and data from 482 blood samples (corresponding to the control and the experiments at 200 mg/kg tetracycline, 500 mg/kg valproate, 300 mg/kg isoniazid, and 2000 mg/kg carbon tetrachloride) and performed principal component analysis (PCA, Fig. 4a). Plotting the trajectory of the 3 rats dosed with 200mg/kg of tetracycline on the 2-dimensional principal component parameter space (using the top 2 principal components PC1 and PC2) reveals highly similar trajectories (colored points in Fig. 4b) that move to the upper right region, before returning to the healthy region in the lower left. The control rats shown in gray remain in the healthy region through the entirety of the 21-day study. The sorted distribution of the linear weights of the 4,302 TVGs for PC1 and PC2 are shown in Fig. 4c. Similar trajectories for observed for isoniazid, valproate, and carbon tetrachloride (Supp. Fig. S19).

**Figure 4.**
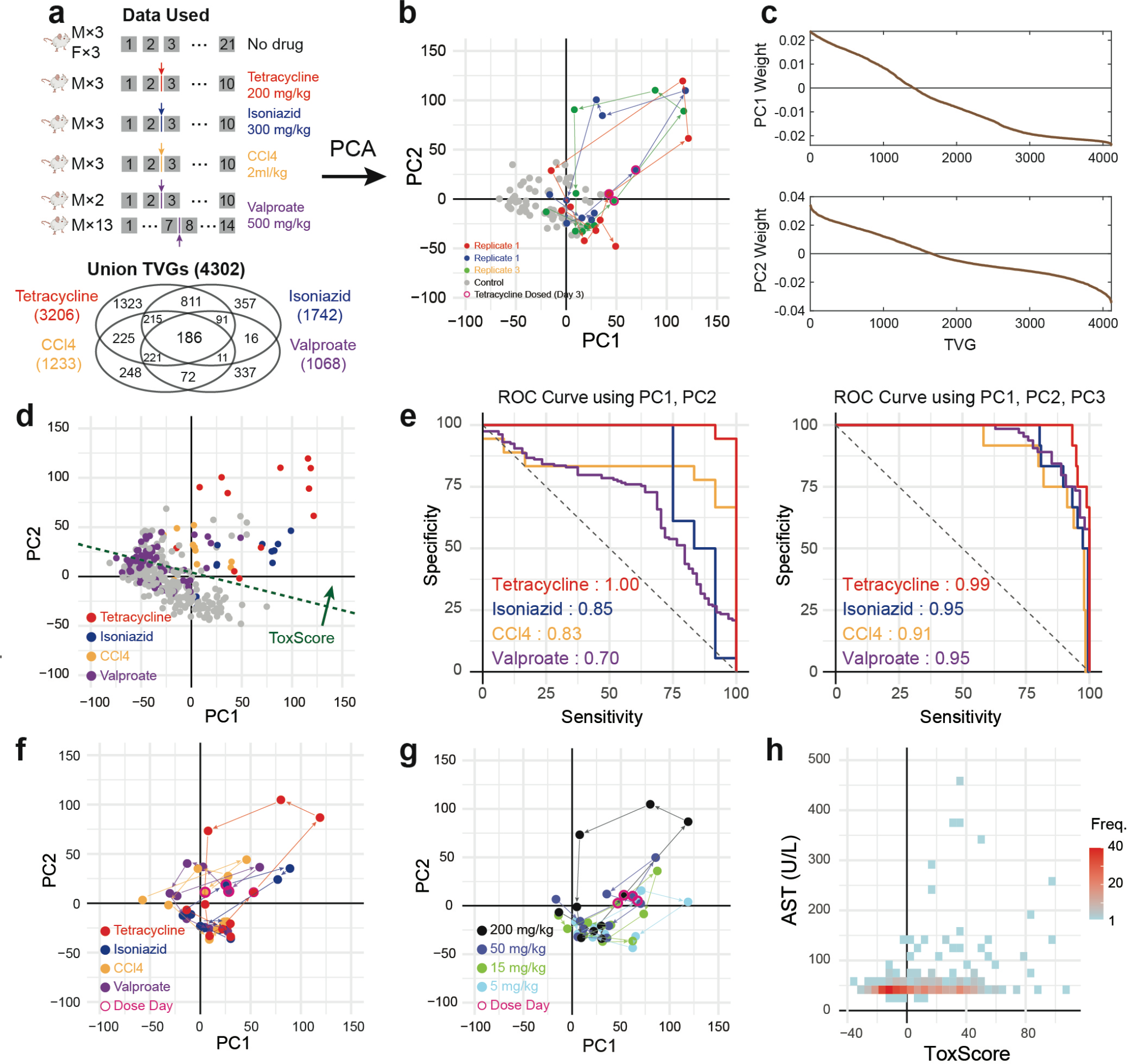
Recovery trajectories in RNAseq principal component space. (a) We performed principal component analysis (PCA) on 4,302 TVGs (the union of TVGs for all 4 drugs), using data from 482 blood samples across a total of 30 rats. In addition to the number of intersection TVGs shown, there were 134 TVGs for tetracycline and valproate only, and 55 TVGs for isoniazid and carbon tetrachloride. The remaining 358 blood samples were not used for PCA because they used lower doses of drugs or resulted in animal death. (b) Trajectories of 3 individual rats dosed with 200mg/kg tetracycline and 6 individual rats not dosed with any drug. All 3 of tetracycline-dosed rats followed a predictable trajectory following dosing, returning to the pre-dosing parameter space after Day 6, indicating recovery. (c) Weights of TVGs for principal components PC1 and PC2, shown in sorted descending order (weights are independently sorted for each PC). (d) Linear separation of Day 4, 5, and 6 TVG expression profiles (colored dots) following high concentration drug dosing vs. control. The perpendicular ToxScore metric indicates perturbation from liver health. (e) Receiver operator characteristic (ROC) curve for each of the 4 drugs, based on varying the ToxScore threshold for calling a positive. 3 of the 4 of the drugs showed area under the ROC (AUROC) values of over 80%, with only valproate at 70% AUROC. Right panel shows that prediction performance is further improved by considering each drug independently using 3 principal components to between 91% and 99%. (f) Recovery trajectories for the different drugs at highest dose. (g) Temporal TVG expression in response to lower tetracycline doses in PC1/PC2 space. (h) Comparison of ToxScore vs. the traditional liver biomarker aspartate transaminase (AST). High AST values is almost always associated with high ToxScore, but the converse is not true. This suggests that ToxScore may have higher sensitivity to liver damage and recovery than AST.

Based on the PC space trajectories, it appears that data from the second and third days after dosing (Days 4 and 5 for most experiments) typically provide the most dramatic expression change relative to control, followed by a moderate expression change in Day 6. Taking expression from Days 4, 5, and 6 as “positives” for drug response perturbation, and all other days as “negatives” for normal health, we find that there is dividing line that separates the positives from the negatives with good sensitivity and specificity. We can represent this separating line as the 0 isocline for a ToxScore that is perpendicular to the separating line:

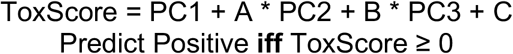

In Fig. 4d, we show the projection of the gene expressions into 2-dimensional PC space using only PC1 and PC2; the green dotted separating line shows A=5, B=0, and C=-50. By varying the intercept C, we can achieve different tradeoffs between sensitivity and specificity, generating a receiver operator curve (ROC) for each of the 4 dosed molecules (Fig. 4e). The area under the ROC curve (AUROC) were moderately high for all 4 molecules (70%, 83%, 85%, and 100%), with valproate exhibiting the lowest AUROC. By expanding the projection to include PC3 and considering each drug molecule independently, we find separating hyperplanes that improves the AUROC to between 91% and 99%. We were pleasantly surprised by this performance, given that the TVG expression data fed into the PCA were not specifically labeled and trained for the task of separating Days 4, 5, 6 from the remaining Days and the control rats. See Supp. Figs. S20-S22 for AUROC results using between 1 and 4 principal components.

For better visualization, Fig. 4f shows the trajectory of the TVG expression levels for the 4 drugs in the two-dimensional PC1/PC2 space. Trajectories for all 4 drugs followed a movement to the upper right, followed by movement back to the lower right. The post-dosing TVG expression levels for rats subjected to lower doses of the molecules provided less separation in parameter space (Fig. 4g, Supp. Fig. S23-S24), consistent with our findings of lowered numbers of TVGs in Fig. 3e.

Because all 4 of the molecules dosed are known to cause liver toxicity, these results suggest that the ToxScore could be potentially used as a universal diagnostic biomarker of liver damage response. To test this hypothesis, we compared the ToxScore vs. the traditional liver biomarker aspartate aminotransferase (AST) values [14] observed for the rats from each blood sample (Fig. 4h). There appears to be a weak correlation between ToxScore and AST, driven primarily by the large number of blood samples with high ToxScore but low AST levels. Additional longitudinal studies dosing animal models with additional liver toxic and non-liver toxic molecules could potentially validate a refined version of ToxScore based on TVGs as a more sensitive liver biomarker than AST. See Supp. Fig. 25 for histopathology results on the rat livers following termination of the experiments.

### Individual Variability in Drug Response

In our first set of experiments dosing the rats with 500mg/kg valproate, 1 of the 3 rats died within one day of dosing and the other two recovered after 1 day and 2 days. The variability in the outcome occurred despite the rats’ similarity in age (10 weeks), weight (220-240g), and sex (male) of the rats. To confirm that this result was not an operator technical error, we performed a follow-up experiment using an additional 15 male rats, all dosed with 500 mg/kg. Additionally, to confirm that minor differences in age were not the cause of the divergent phenotypic results, we selected 3 different ages for the rats: 3-4 weeks, 8 weeks, and 8-9 months (Fig. 5a). The outcomes of the experiments confirmed the individual variability in the outcomes (Fig. 5b): 2/15 rats died 1 and 2 days after valproate injection (Death Group), 9/15 rats showed phenotypic changes on Day 8 (Slow Recovery group), and 4/15 rats exhibited no symptoms (Fast Recovery group).

**Figure 5.**
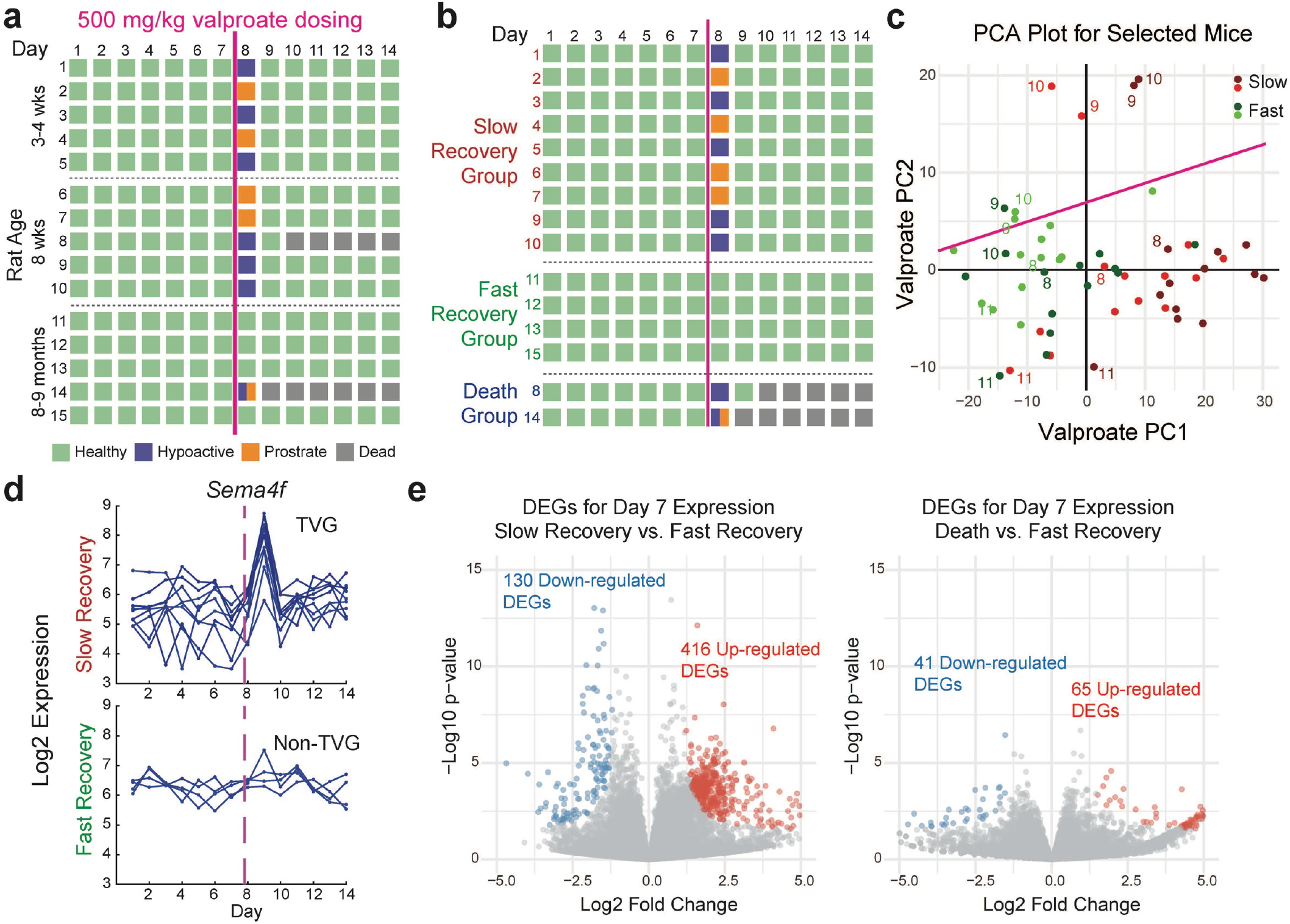
Expansion study on individual outcome variability for rats dosed with 500 mg/kg valproate. (a) Experimental design and phenotypic outcomes. (b) Animal experiments sorted by outcome phenotype into Fast Recovery, Slow Recovery, and Death groups. (c) Expression trajectories on parameter space constructed via PCA on the 1,068 valproate TVGs. Expression profiles of Slow Recovery rats deviated significantly farther on Days 9 and 10 from the lower healthy cluster than the Fast Recovery rats. (d) Example TVG with larger response in the Slow Recovery group than the Fast Recovery group. (e) DEGs identified based on Day 7 expression profiles comparing Slow Recovery to Fast Recovery group (left) and Death to Fast Recovery group (right).

Given these results, we next asked whether the differences in phenotypic outcomes could be recapitulated through the valproate TVG genes. For ease of data analysis, we performed a new PCA only on the 1,068 valproate TVGs, and obtained new PC1 and PC2 weights only on these genes. The expression trajectories of the Slow Recovery group deviated significantly farther on Day 9 and Day 10 from the healthy region of the PC1/PC2 parameter space, compared to the Fast Recovery group. This result suggests that the phenotypic symptoms can be effectively captured through blood TVG expression (see Supp. Fig. S26 for additional trajectories). Fig. 5d shows the different temporal expression profiles of the *Sema4f* gene for the Fast Recovery and Slow Recovery groups.

The Fast Recovery and Slow Recovery animals appeared to in slightly different subregions of the healthy region in PC1/PC2 parameter space. To further analyze potential pre-dosing gene expression differences that may be prognostic of phenotypic outcomes, we performed DESeq2 analysis on the full RNA expression profiles from Day 7, just before the animals were dosed (Fig. 5e). We identified 546 DEGs comparing the Slow Recovery and Fast Recovery groups and 106 DEGs comparing the Death and Fast Recovery groups (see Supp. Fig. S27 for list of DEGs and pathway analysis). We wish to caveat the reader that we cannot be sure that the DEGs identified are not reflecting age-based differences in expression, because all of the Fast Recovery animals happened to be from the 8-9 month age group.

## Discussion

Through the course of this set of experiments, we collected roughly 5Tb of Fastq NGS data from 829 RNAseq samples. To facilitate analysis of this large longitudinal RNA expression data, we developed and utilized several ad hoc bioinformatics methods to (1) minimize the number of false positive DEGs called from the large number of potential pairwise differential expression analyses, and (2) identify TVGs from longitudinal data. Although these tools have not be rigorously optimized for performance, spot check evaluation results suggests that the TVG calling is reasonably accurate and catches a significant number of false positive DEGs called based on cross-sectional tools.

We made several qualitatively unexpected findings: First, even in the control group, there were roughly 300 TVGs, and many of these were part of what appeared to be an oscillatory expression pattern with a 2 week period. Second, many TVGs were activated or repressed at different drug dose thresholds, suggesting that rat biology include multiple layers of response mechanisms activated at different levels of toxicity severity. Third, the relatively small number of genes that were called as TVGs for all 4 drugs suggests that there may be multiple separate recovery/detoxication mechanism biologically in rats. Finally, the same drug dose can elicit very different phenotypic outcomes (including Death, Slow Recovery, and Fast Recovery) in different individuals, and these phenotypic outcomes could be reflected in the degree and duration of response in TVGs. Importantly, all of these observations required high frequency longitudinal data, and are not visible from a single before/after comparison. From the findings in this work, <u>we advocate for broader adoption of high frequency longitudinal RNA expression profiling</u>.

In addition to biological findings, our results also suggest the strong potential of longitudinal RNAseq to serve as molecular diagnostic tests. Our ToxScore metric, a linear combination of the top 3 principal components from PCA provided drug response AUROC values between 91% and 99% for the 4 drug molecules. One immediate application may be in the prognosis of individualized patient responses to drugs with known liver toxicity, in order to inform personalized therapy selection or dosage. One potential complication in realizing this goal would be the logistical complexity of collecting high frequency blood samples from human patients on a daily basis.

Longitudinal RNA expression studies can be costly to conduct because the large number of RNAseq libraries that need to be constructed and sequenced. In the experiments presented here, we typically would take 10 or more daily blood samples per animal, corresponding to 5x more samples than a simple before/after dosing analysis. New technologies and methods for improving the total cost of longitudinal RNAseq studies would be greatly synergistic with and beneficial to the adoption of longitudinal RNAseq-based diagnostics. Cost reduction would need to be pursued in every step of the process, including animal handling, blood collection, RNA extraction, NGS library preparation, NGS chemistry, and data analysis.

The data analysis methods we used in this work are primarily human-driven, using traditional tools based on linear combinations of gene expression levels. Given the depth and complexity of RNA expression data, it is reasonable to assume that a well-trained deep neural network would be able to better utilize the “long tail” of information inherent within gene expression variations that are below our stringent cutoffs needed to avoid false positives. Deep neural networks [15], such as those based on transformers [16], require a large amount of data to achieve performance significantly superior to traditional machine learning models. In our dataset of 829 samples, roughly 9,000 genes were expressed at non-zero levels in each sample, so the total size of this dataset is roughly 7.56 million tokens. 7.46M tokens is large for biological datasets, but just a small early step in comparison to large language models (LLMs). For example, GPT-1 [17] was trained on the BooksCorpus dataset [18], corresponding to approximately 250M tokens. Given the remarkable emergent properties of LLMs as they are provided more parameters, floating point operations, and training data, we look forward to the impact of domain-specific biology AI.

Our dataset is publicly available for download from NCBI SRA and NCBI GEO at https://www.ncbi.nlm.nih.gov/bioproject/PRJNA1026523 (PRJNA1026523). Because this dataset is the first high frequency longitudinal RNAseq dataset, encapsulating 4x as many samples as the previous largest rat RNAseq studies on the GEO database [19], we believe that deeper analysis than what we have performed in this work could yield additional qualitative and quantitative findings. We encourage interested readers to independently analyze and report findings from the data, and welcome academic collaborations as we continue to collect larger scale longitudinal omics data on animal models.

## Acknowledgements

The authors thank Nikolay Bazhanov, Alessandro Pinto, and Sherry Chen for assistance and advice on experimental design and data analysis. No federal research funds were used for these studies.

## Author Contributions

AG and DYZ conceptualized the study and directed the research. WC, YC, EYM, and DYZ designed the experiments. WC, YC, and QJ performed bioinformatic pre-processing. WC, YC, and DYZ created and optimized metrics for identifying DEGs and TVGs. WC, YC, and DYZ wrote the paper with input from all authors.

