## Supplemental File for "High Frequency Longitudinal RNAseq Reveals Temporally Varying Genes and Recovery Trajectories in Rats"

**Supplementary Information**

Wei Chen[1,2], Yi Chai[1,2], Qi Jiang[1], Eva Y. Miao[1], Ashwin Gopinath[1]*, David Yu Zhang[1]*
[1] Biostate AI, Inc., [2] Equal Contributors

### Materials & Methods

#### 1. Chemicals

Saline solution (Cat# R11B23063006) was purchased from Shandong Qidu Pharmaceutical (China). TRIzol (Cat# Abs60154) was purchased from Aibixin Biotechnology (China). Chloroform (Cat# 10006818) was purchased from Sinopharm Chemical Reagent Company (China). Ficoll-Paque PLUS (Cat# 17144002) was purchased from Cytiva Life Sciences (USA). Isoniazid (Cat# HY-B0329) was purchased from MedChemExpress (China). Valproic acid (Cat# V298968-100ml) and Tetracycline hydrochloride (Cat# T140622-100g) were purchased from Aladdin Scientific (China). Carbon tetrachloride (Cat# 10006464) were purchased from Sinopharm Chemical Reagent (China).

#### 2. Animals and animal care

Animal experiments in this study were approved by the Animal Care Committee at Biostate.AI (approval number: 101) and HD Biosciences Co. Ltd (approval number: 118-8). Sprague-Dawley rats were purchased from certified provider - Charles River Laboratories International, Inc (Shanghai, China). Rats were acclimatized prior to the experiment and were housed in enriched and ventilated housing cages throughout the experimental phase. Cage litter was changed at least once a week. The rats were housed under specific-pathogen-free (SPF) condition and subjected to a normal 12 hours light and night cycle, at 22 ± 2°C and 50 ± 10 % relative humidity. Chow and water were available *ad libitum*. The health conditions were examined and recorded daily by a veterinarian. Detailed examination contents are shown in Table S1.

#### 3. Blood samples and liver samples collection and storage

250 μL of whole blood was collected daily from the same site of jugular vein of each rat between 10:00 am and 10:37 am. Whole blood samples were immediately processed for RNA extraction and liver function biomarker testing. On the final day of the study, rats were fasted overnight and then euthanized by overdosing with pentobarbital (120 mg/kg) through intraperitoneal injections. Whole blood and liver samples were immediately collected under RNase-free conditions. Whole blood samples were obtained via heart puncture. 1 mL of whole blood underwent the standard PBMC separation process, and the rest was preserved at -80°C for RNA extraction. Liver samples were preserved at -80°C and used for RNA extraction and histological analysis.

#### 4. RNA extraction from whole blood samples and liver samples

For RNA extraction from whole blood, 100 uL of whole blood was mixed with 700 uL of Trizol reagent and then underwent standard TRIzol and Chloroform RNA extraction method. Extracted RNA was further purified using Automatic Nucleotide Isolation Machine (Cat# NPA-32P, Bioer Technology, China) with MagaBio Plus Total RNA Purification Kit (Cat# BSC53M1B, Bioer Technology, China).

For RNA extraction from liver samples, around 30 mg of chopped liver tissue was mixed with 700 uL of TriZol Reagent and Lysing MatrixD. The mixture was ground for 2 min and then mixed with 140 uL of Chloroform. After 2 min of incubation at room temperature and 10 min of centrifugation at 12000 rpm and 4°C, supernatant was transferred to a tube and was further purified with MagaBio Plus Total RNA Purification Kit.

Purified RNA was quantified by using Nanodrop 2000 (Thermo Fisher, USA). The quality of RNA was measured by using RNA ScreenTape Assay (Agilent, USA) and 4200 TapeStation System (Agilent, USA).

### 5. mRNA capture and Release

VAHTS mRNA Capture Beads (Cat# N401-02, Vazyme Biotech, China) was used for the enrichment of mRNA from total RNA extracted from blood and tissue samples. During the mRNA capture and release process, mRNA was fragmented.

### 6. cDNA Synthesis and Library Preparation

Library preparation of mRNA including three steps: reverse transcription, adaptor ligation, index PCR amplification. Reverse transcription and adaptor ligation were performed using VAHTS Universal V6 RNA-seq Library Prep Kit for Illumina (Cat# N604-2, Vazyme Biotech, China) with standard protocol. Index PCR amplification was performed using VAHTS RNA Multiplex Oligos Set 1 for Illumina (Cat# N323-01, Vazyme Biotech, China) and VAHTS RNA Multiplex Oligos Set 2 for Illumina (Cat# N324-01, Vazyme Biotech, China) with standard protocol. One round of 1.6 × beads (Cat# CNGS-0500, CleanNA, Netherlands) purification was then performed after adaptor ligation. One round of 1.6 × beads purification was performed after index PCR amplification. Library DNA were quantified by Qubit 4.0 using Equalbit 1 × dsDNA HS Assay Kit (Cat# EQ121-01, Vazyme Biotech, China). Library was analyzed by using D1000 ScreenTape Assay (Agilent, USA) and 4200 TapeStation System.

### 7. Sequencing

2 ×150 paired-end sequencing was performed on Illumina NovaSeq 6000 system (Illumina, USA) at Sequanta (China).

### 8. Liver function Analysis

Liver function tests of 4 liver function related biomarkers - alanine transaminase (ALT), aspartate aminotransferase (AST), alkaline phosphatase (ALP) and gamma-glutamyl transferase (GGT) were performed on Hitachi Automatic Analyzer 3500 (Hitachi, Japan) following IFCC methods.

### 9. Animal Study - Baseline Study – Bleeding related gene investigation

3 male and 3 female Sprague-Dawley rats (SD Rats) were acclimatized for 21 days with daily 0.25 mL blood extraction and without any substances administration. Detailed study design is shown in Supplementary Excel 3.

### 10. Animal Study - Toxic Study

39 male SD rats were acclimatized for 10 days with daily 0.25 mL blood extraction. Rats in the Gcon group (control) were intraperitoneally injected with 0.9% sodium chloride on day 3. Rats in the GA group were administered a 1:1 mixture of corn or olive oil and carbon tetrachloride orally at different dosages (0.5, 1, 1.5, 2 ml/kg) on day 3. Rats in the GB group were intraperitoneally injected with Isoniazid dissolved in sterile saline at different dosages (400, 800, 1600, 3200 mg/kg) on day 3. Rats in the GC group were intraperitoneally injected

with Valproic acid diluted in sterile saline at different dosages (250, 500, 1000, 2000 mg/kg) on day 3. Detailed study design is shown in Supplementary Excel 3.

### 11. Animal Study - Sub Toxic Study

24 male SD rats were acclimatized for 10 days with daily blood extraction. Rats in GB group were handled as described above but with different dosages (10, 30, 100, 300 mg/kg) on day 3. Rats in GD group were intraperitoneally injected with tetracycline hydrochloride dissolved in sterile saline at different dosages (4, 15, 50, 200 mg/kg) on day 3. Detailed study design is shown in Supplementary Excel 3.

### 12. Animal Study - Age investigation

15 male SD rats were acclimatized for 14 days with daily blood extraction. All rats were intraperitoneally injected with Valproate at dosage of 500 mg/kg on day 8. In GC5 group, the rats were 3-4 weeks old. In GC6 group, the rats were 8 weeks old. In GC7 group, the rats were 8-9 months old. Detailed study design is shown in Supplementary Excel 3.

### 13. Adaptor Trimming, Alignment and Normalization

The FASTQ file was initially trimmed to remove Illumina adaptor sequences, and subsequently aligned with the Rattus norvegicus reference genome (NCBI GCF_015227675.2, Genome assembly mRatBN7.2) using HISAT2 (default parameters). This process generated a raw gene expression file. To minimize differences in sequence depth across various samples, six steps of normalization were performed:
1. Identification of genes with raw read counts greater than 0 across all samples.
2. Transformation of data using the log2 method.
3. Calculation of the mean log2 expression for each gene across all samples.
4. Subtraction of the log2 expression of each gene in each sample from the mean values, resulting in Δ expression.
5. Calculation of the median Δ expression (sample factor) for each sample across all genes.
6. Adjustment of the log2 expression of each gene in each sample by subtracting the corresponding median Δ expression values.

### 14. DEG, TVG, and pathway enrichment analysis

We used DESeq2 for performing fundamental paired DEG analysis on the raw count with 0 or low expressed genes (sum of count across samples < 10) filtered out, with cutoff $|\mathrm{log2FoldChange})| > 1, \mathrm{Padj} < 0.05$ in most of the analysis. To reduce false discovery rate, we applied one more criterion: $-\log 10(pvalue) + \log 10(0.05) > \frac{1}{|log2FoldChange|-1}$. On the base of that, we identify TVGs by the sum of significance score calculated in each comparison as score = $\min(-log10(padj) - (-log10(pcutoff = 0.05), 20)$, where score for each comparison are capped at 20 to prevent overweight single comparison and better capturing the time-course expression difference. TVGs are then ranked and eyeballed to find the suitable cutoff based on one gene's expression fluctuation. Pathway enrichment analysis are performed using R package cluserProfiler (https://bioconductor.org/packages/release/bioc/html/clusterProfiler.html) based on rat genome, and statistical cutoff of Benjamini-Hochberg (BH) corrected p-value < 0.05.

### 15. PCA and trajectory analysis

For PCA analysis, we used the unions of TVGs identified from each drug, excluding 300 bleeding TVGs and low count (sum up < 10 across all samples) genes. Variance stabilizing transformation (VST) was applied to stabilize variance across the dataset for PCA analysis. We chose days immediately upon drug injections as true positive (day 3-6 for isoniazid, CCL4, Tetracyclines and day 8-11 for Valproate), and days of initial or recovery stages (1-2 and 7-10 for isoniazid, CCL4, Tetracyclines; and day 1-7, 12-14 for valproate) a true negative. We first identified the best-separating line that achieved the highest sensitivity (0.855;) and specificity (0.880) across all 4 drugs (PC2 = -0.2 PC1 + 10) and used it for calculating AUC-ROC for each drug respectively. For evaluation on 3 or more dimensions PCs (Supp. Fig. 20-21), logistic regression with the generalized linear model is performed using different combinations of PCs as predictors to maximize the probability of each datapoints belonging to its true classification for each drug respectively, with formula: $Maximize \left\{logit\left(P(Y = true_{status})\right) = \beta 0 + \sum_{i+1}^{n} \beta i \cdot PCi\right\}$

to find the space separating true positive vs negative best. Eventually, we obtained a set of coefficients best separating the two classes pre-defined for each drug, with coefficients documented in Table 1, which is then used for calculating ROC.

### Supplementary Figures

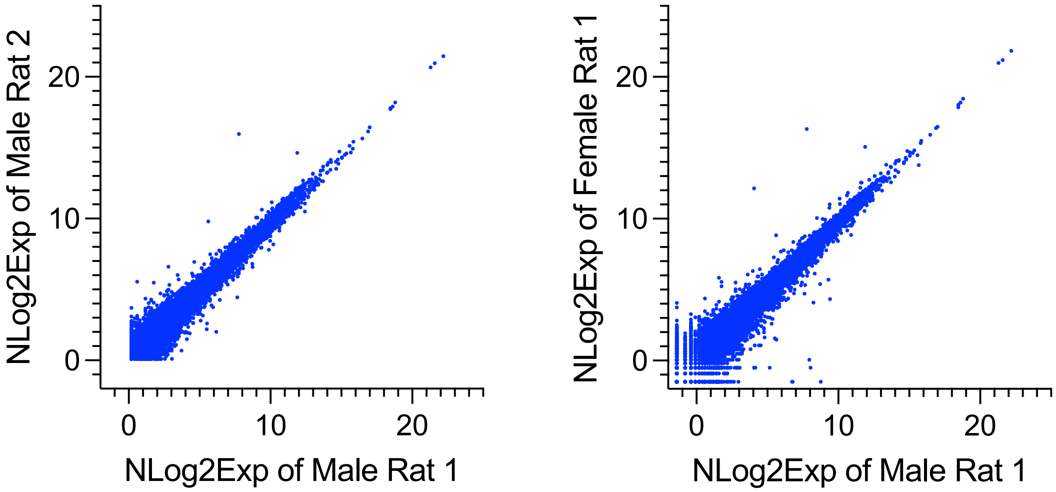


**Fig. S1.** RNAseq reproducibility. Orthogonal plot of normalized log2 gene expression values of male rat 1, male rat 2 and female rat 1.

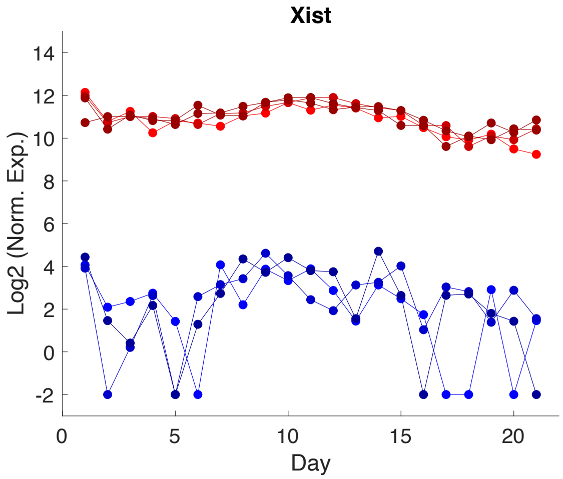


**Fig. S2.** Example longitudinal gene expression profiles for 3 male rats (blue dots) and 3 female rats (red dots) for the sex-based differentially expressed gene Xist.

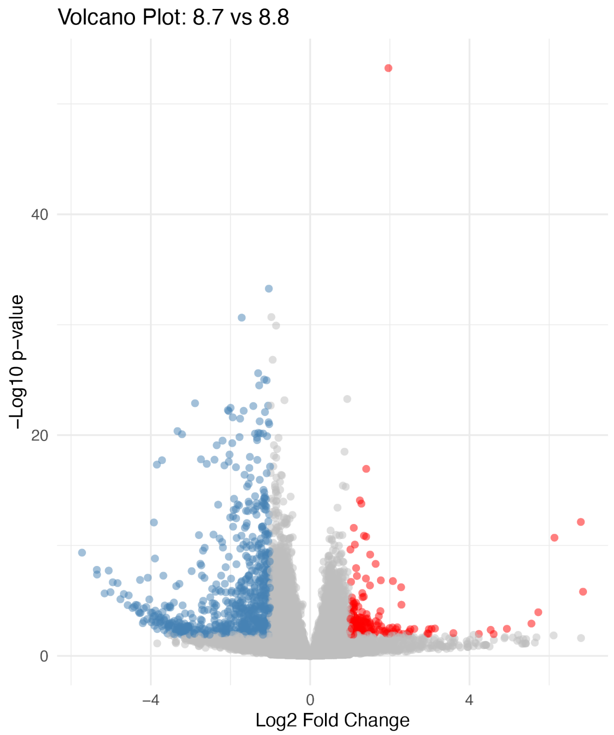


**Fig. S3.** Analysis of differentially expressed genes (DEGs) using DESeq2, comparing Day 2 vs. Day 1 for N = 6 rats. A total of 234 up-regulated and 701 down-regulated DEGs were identified, based on the criteria of $p < 0.05$ and Abs(log2fold) > 1.

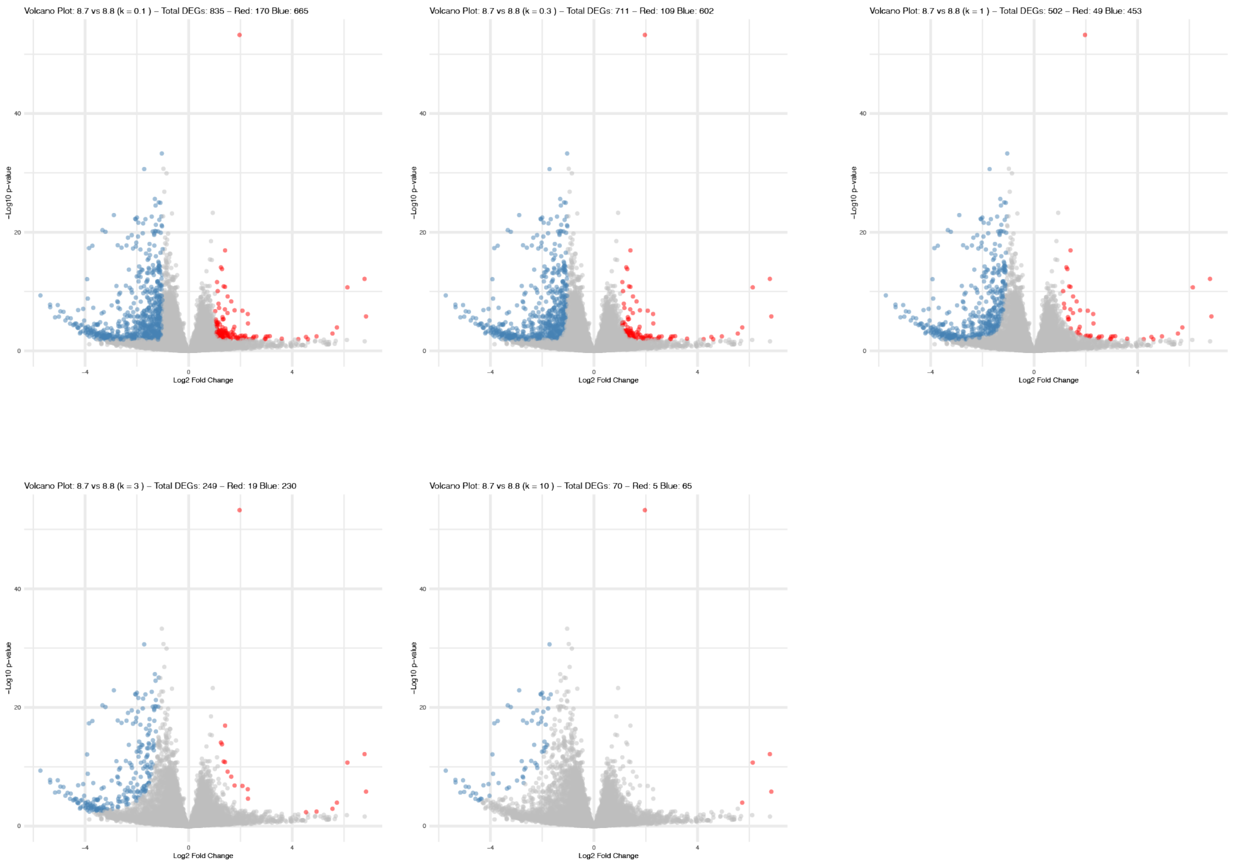

**Fig. S4.** Analysis of differentially expressed genes (DEGs) using DESeq2, comparing Day 2 vs. Day 1 for N = 6 rats. Based on DEG **iff** $[\mathrm{Abs}(\mathrm{log2fold})-1] * \mathrm{Max}\{[-\log(\mathrm{pval}) + \log(0.05)], 0\} \geq k$, k = 0.1, 0.3, 1, 3, 10 criteria.

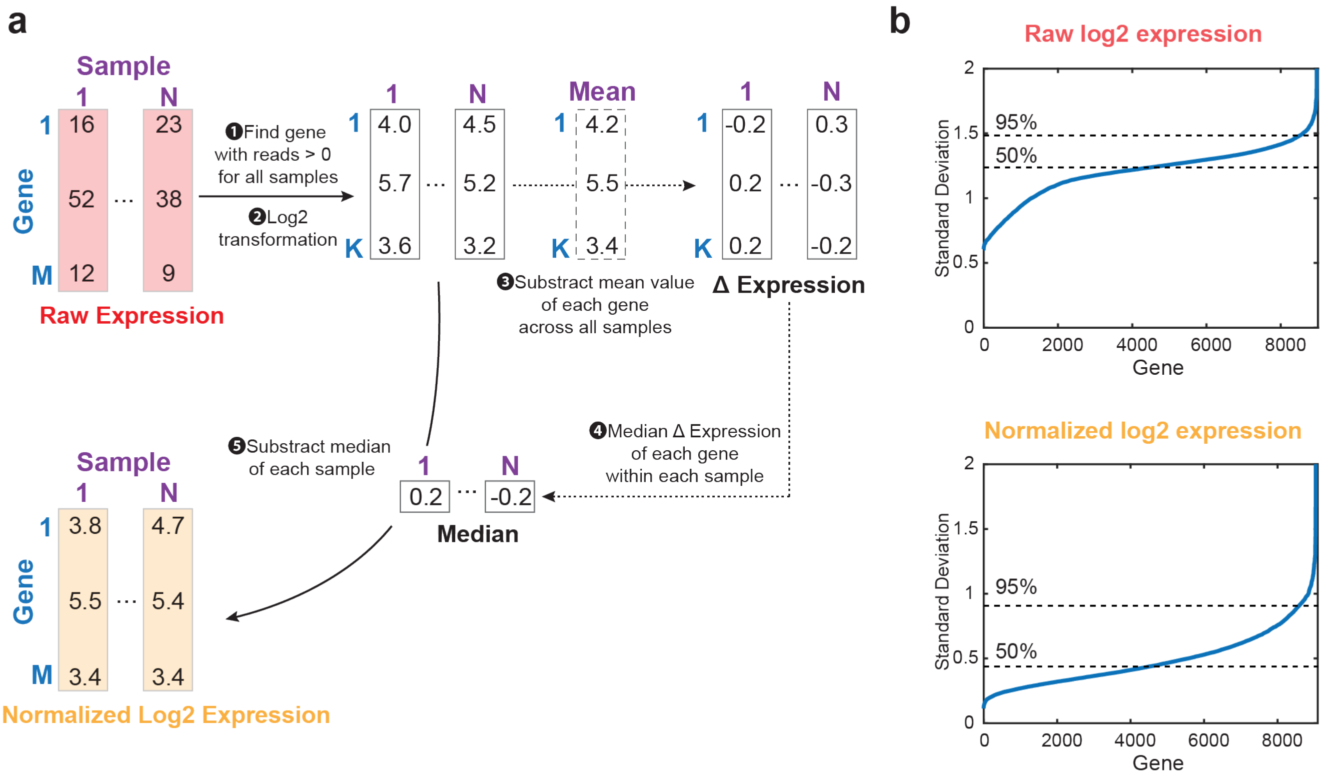


**Fig. S5.** (a) Schematic illustration of raw expression normalization process. (b) Standard deviation of log2 expression of N = 6 rats across 21 days in the control study. Raw log2 expression and normalized log2 expression were plotted respectively.

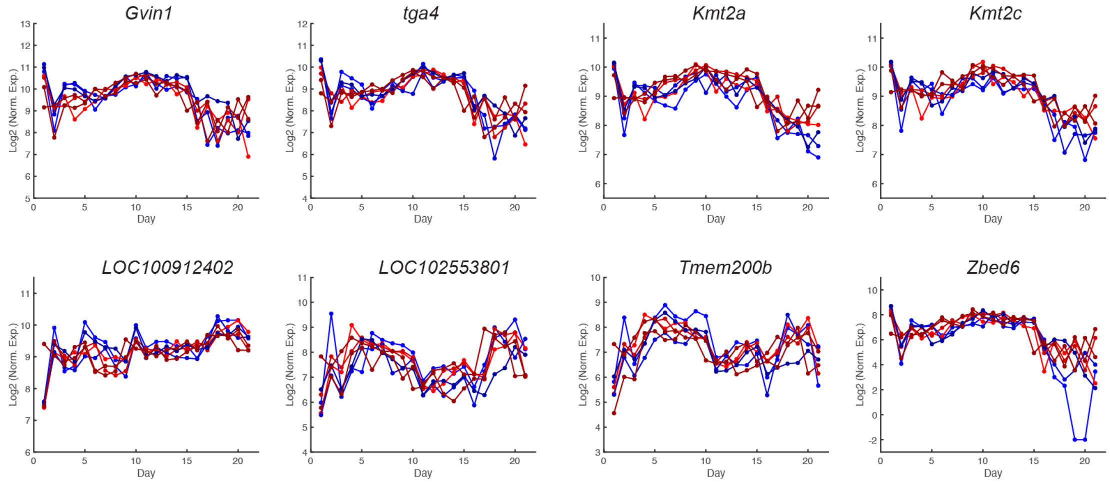


**Fig. S6.** Examples of gene expression with TVG+ and DEG(2/1) + in control studies.

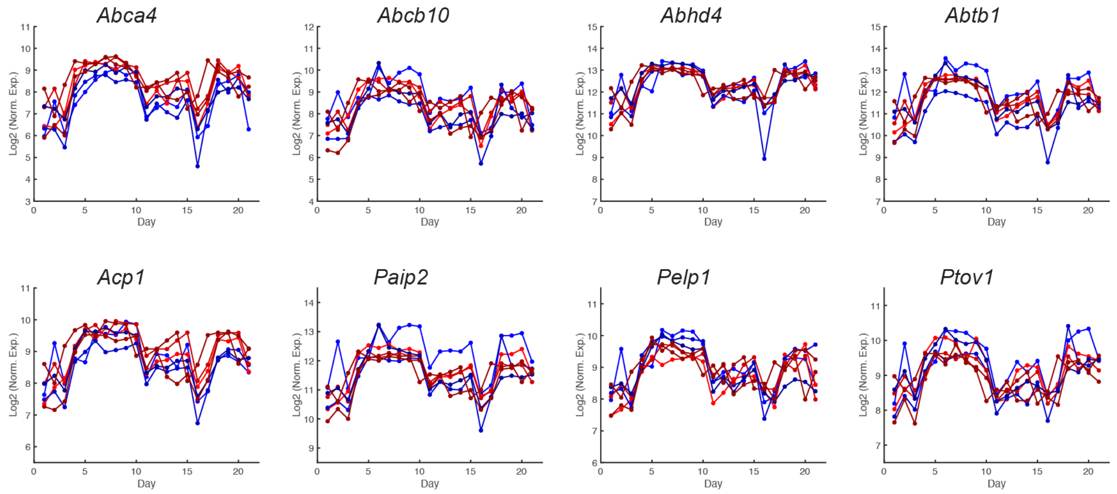


**Fig. S7.** Examples of gene expression with TVG+ and DEG(2/1) – in control studies.

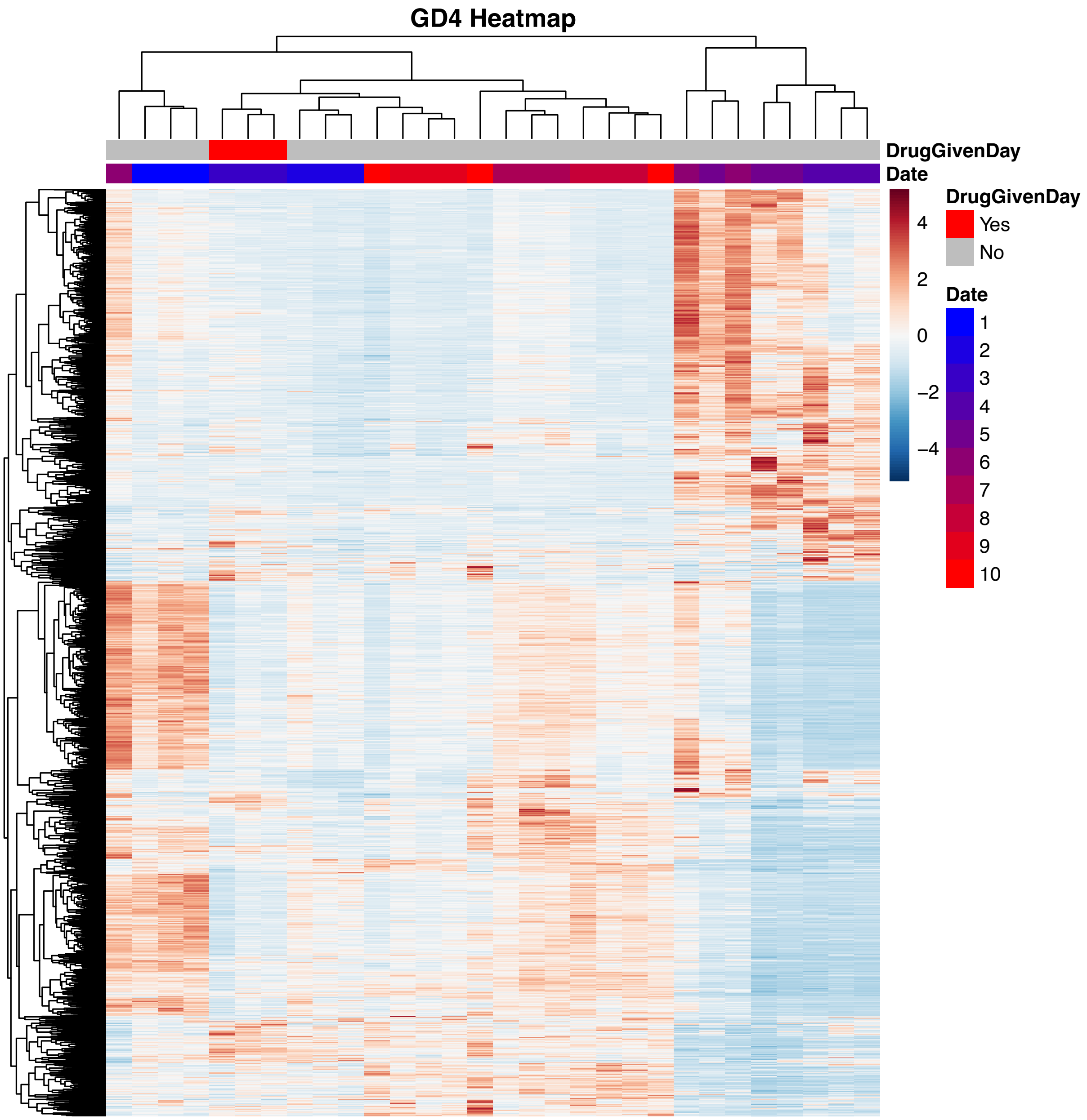

**Fig. S8.** Unsupervised clustering of tetracycline TVGs. Z-Score Normalized RPM Counts excluding globin and bleeding-related TVGs are used for performing unsupervised clustering using Euclidean distances. Genes with low count (sum up < 10 across all samples) are also filtered out and 3202 kept. Column represent daily collected samples of rats injected with 200mg/kg tetracycline. Rows represent z-score normalized count for each gene across samples with red indicating drug administration days and gray for non-administration. The expression of each gene is scaled across samples. Color intensity reflects gene expression deviation from the mean.

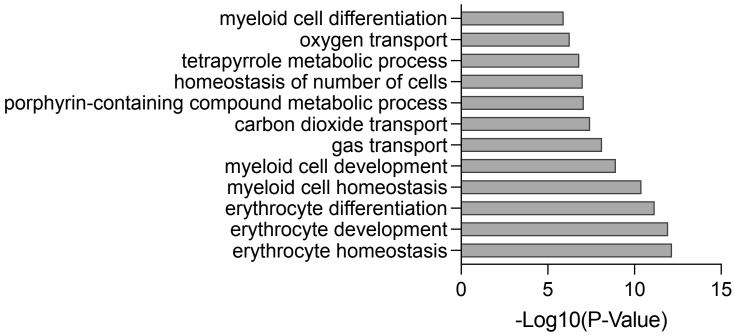


**Fig. S9.** Gene pathway association for Bleeding TVGs from WO1.

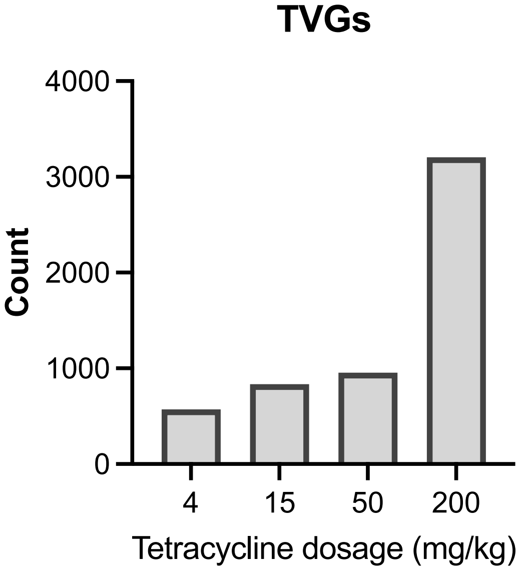


**Fig. S10.** - Statistics and macro analysis of threshold response genes and linear response genes. Additional examples.

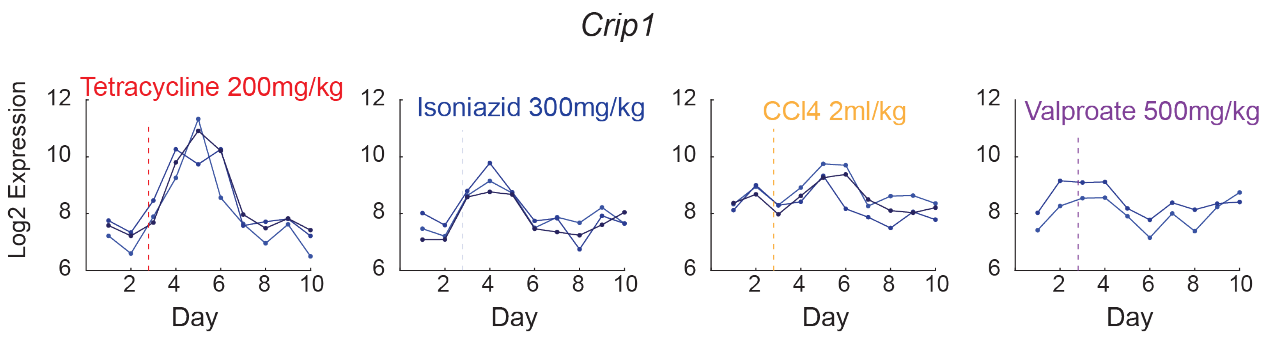


**Fig. S11.** Examples of genes that TVGs for different subsets of drugs.

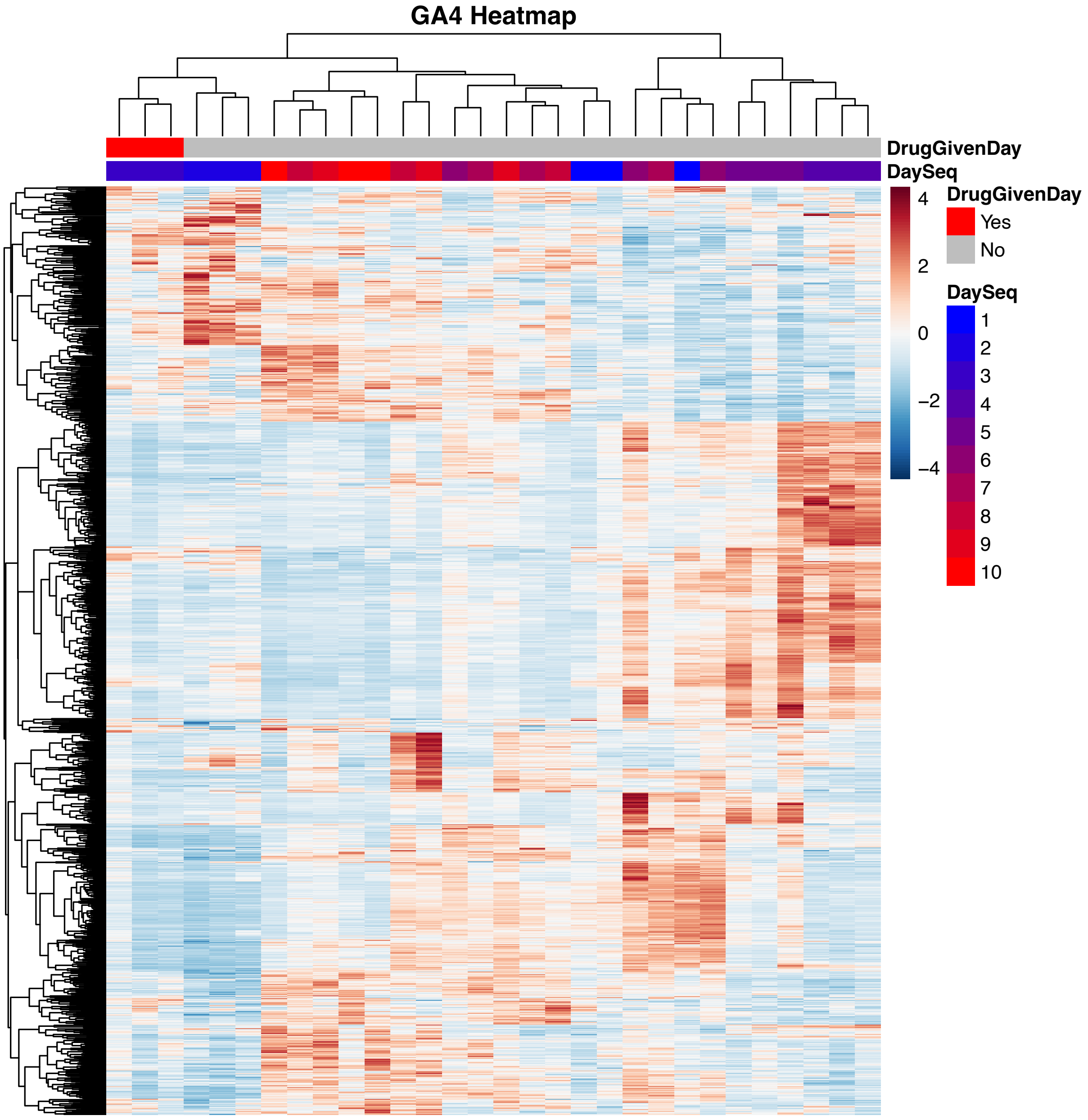


**Fig. S12.** Unsupervised clustering of CCL4 TVGs. Z-Score Normalized RPM Counts excluding globin and bleeding-related TVGs are used for performing unsupervised clustering using Euclidean distances. Genes with low count (sum up < 10 across all samples) are also filtered out and 1246 kept. Columns represent daily collected samples of rats injected with 2ml/kg CCL4. Rows represent z-score normalized count for each gene across samples with red indicating drug administration days and gray for non-administration. The expression of each gene is scaled across samples. Color intensity reflects gene expression deviation from the mean.

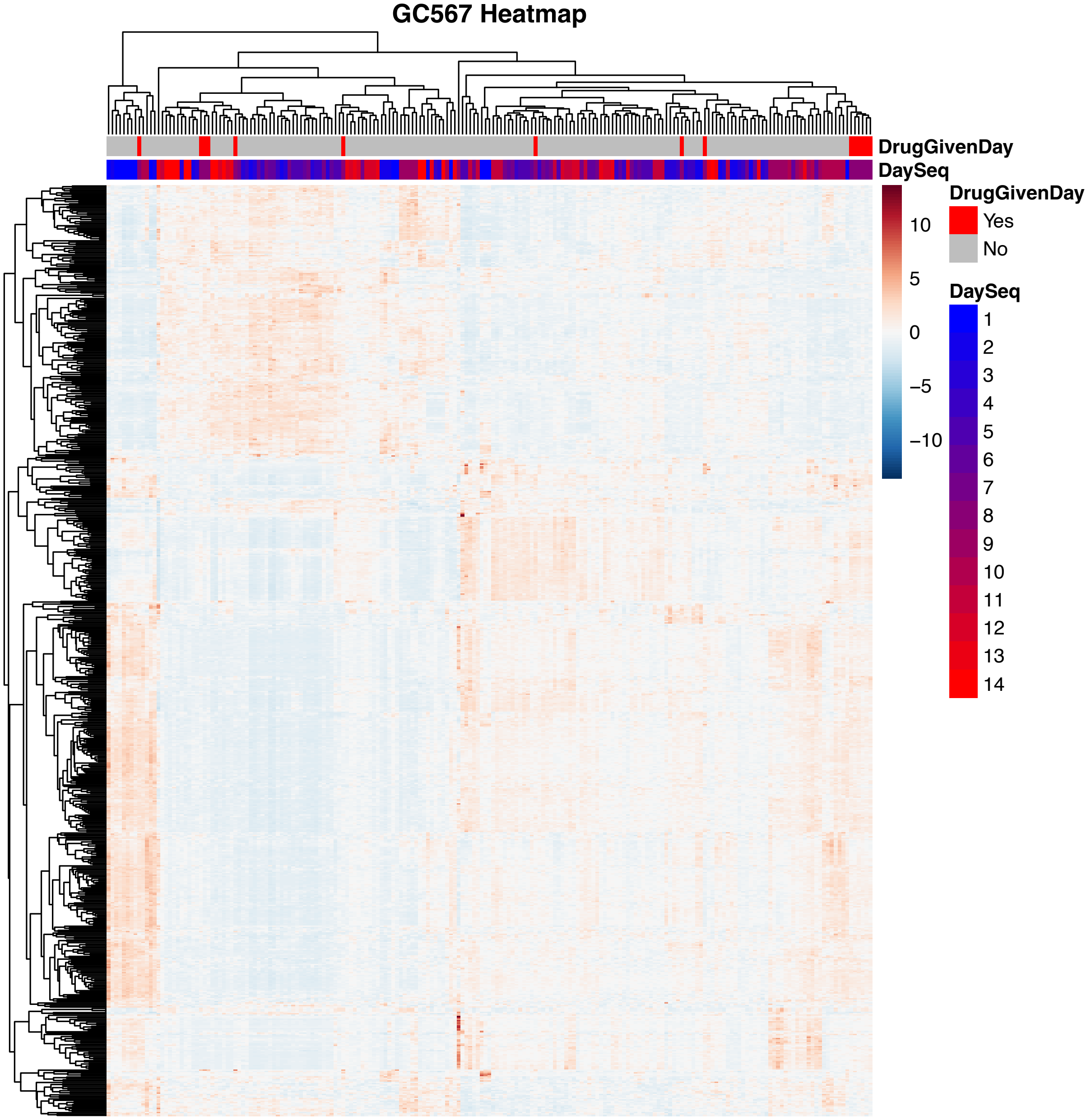


**Fig. S13.** Unsupervised clustering of Valproate TVGs. Z-Score Normalized RPM Counts excluding globin and bleeding-related TVGs are used for performing unsupervised clustering using Euclidean distances. Genes with low count (sum up < 10 across all samples) are also filtered out, and 680 kept. Columns represent daily collected samples of rats injected with 500mg/kg CCL4. Rows represent z-score normalized count for each gene across samples with red indicating drug administration days and gray for non-administration. The expression of each gene is scaled across samples. Color intensity reflects gene expression deviation from the mean.

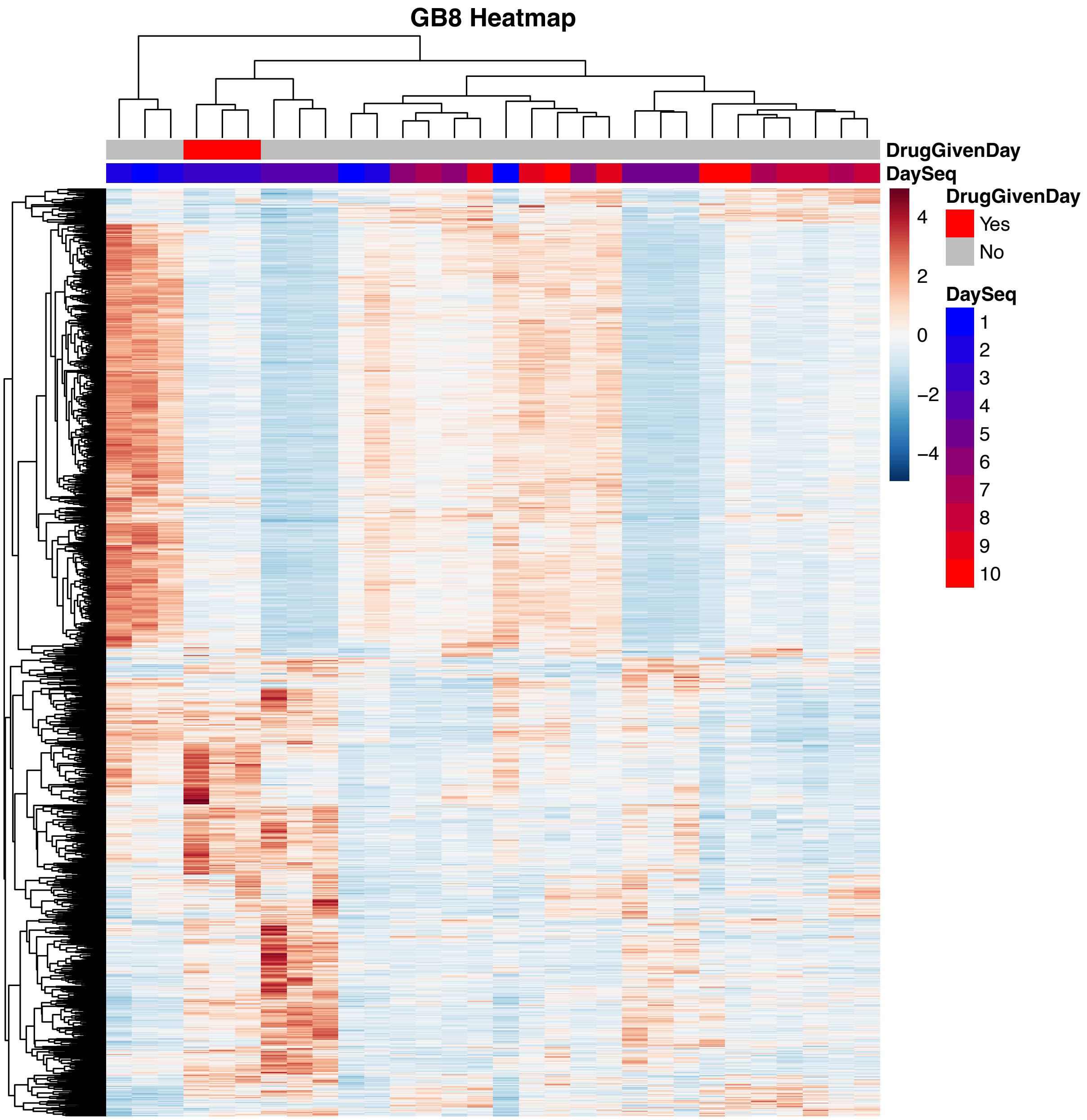


**Fig. S14.** Unsupervised clustering of isoniazid TVGs. Z-Score Normalized RPM Counts excluding globin and bleeding-related TVGs are used for performing unsupervised clustering using Euclidean distances. Column represent daily collected samples of rats injected with 300mg/kg isoniazid. Genes with low count (sum up < 10 across all samples) are also filtered out and 1739 kept. Rows represent z-score normalized count for each gene with red indicating drug administration days and gray for non-administration. The expression of each gene is scaled across samples. Color intensity reflects gene expression deviation from the mean.

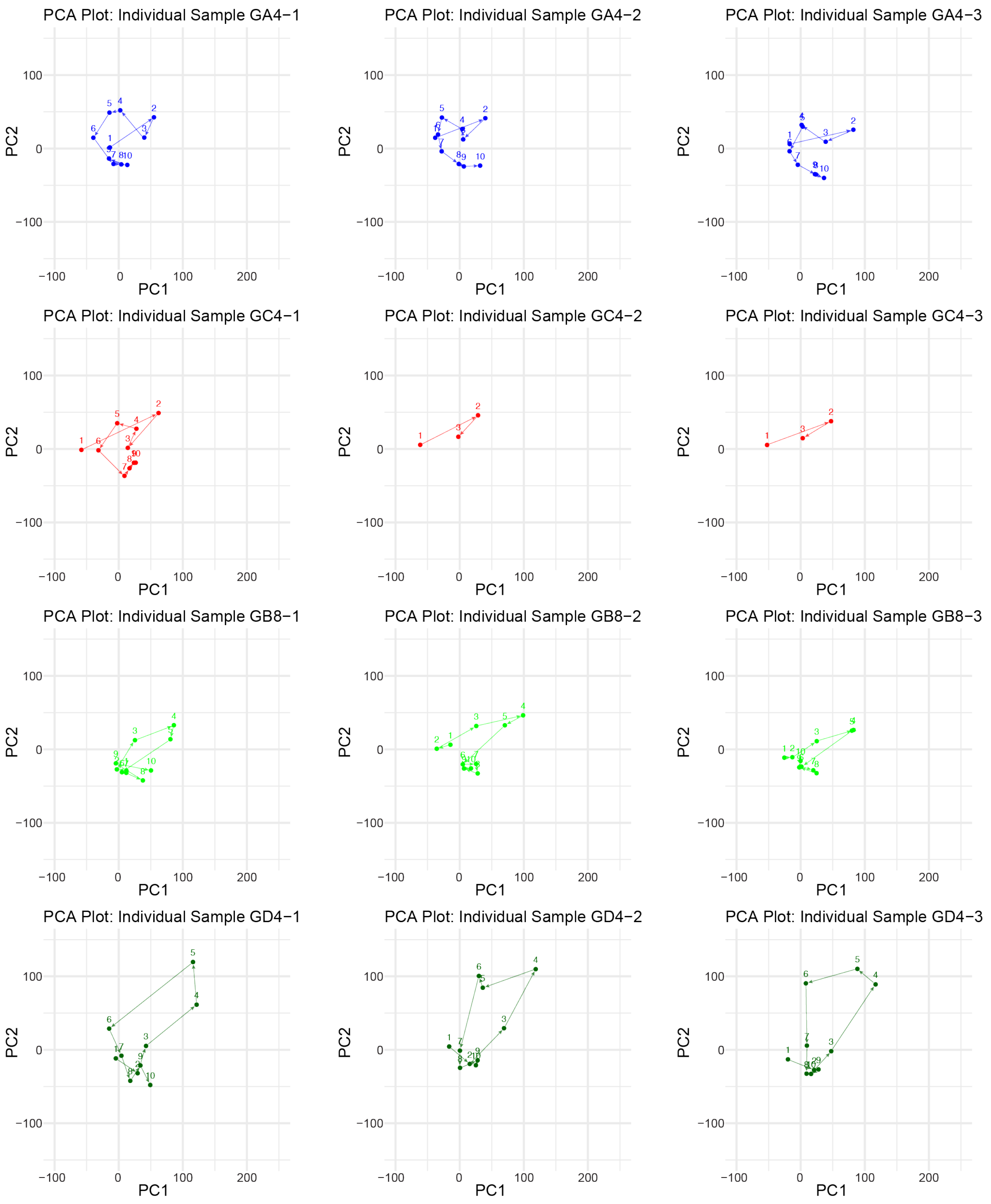


**Fig. S15**. Individual trajectories for highest doses of isoniazid, CCl4, and valproate

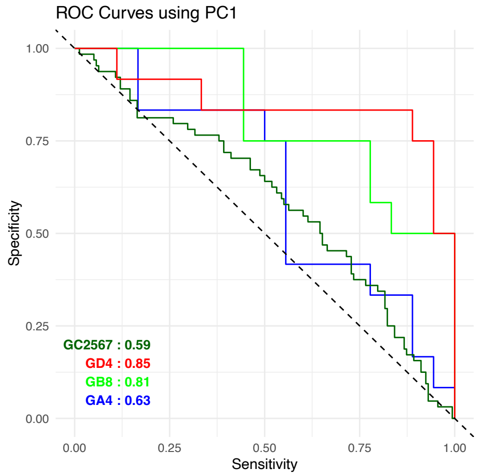


**Fig. S16.** ROC plot for all 4 drugs, using only PC1

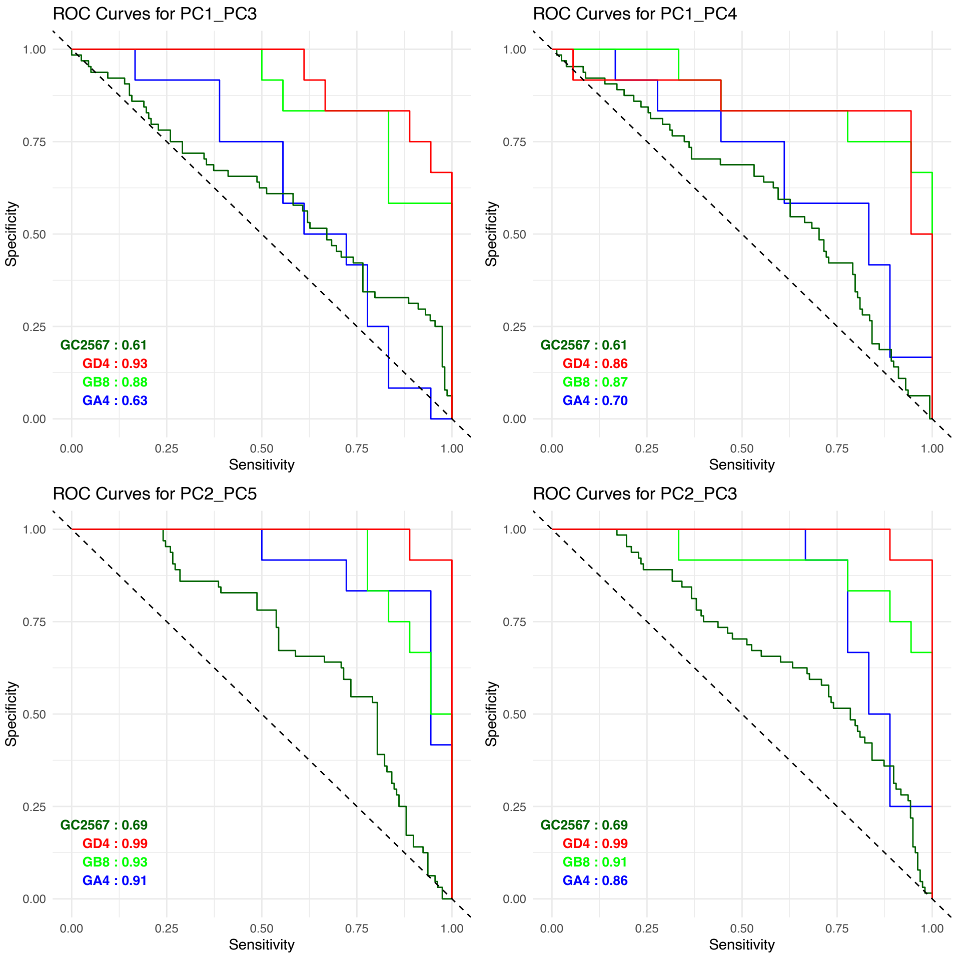


**Fig. S17.** ROC plot for all 4 drugs, using PC1/PC3, PC2/PC3, PC1/PC4, PC2/PC5. For PCA analysis, we used the unions of TVGs identified from each drug, excluding 300 bleeding TVGs and low count (sum up < 10 across all samples) genes. Variance stabilizing transformation (VST) was applied to stabilize variance across the dataset for PCA analysis. We chose days upon drug injections as true positive (day 3-6 for isoniazid, CCL4, Tetracyclines and day 8-11 for Valproate), and days of initial or recovery stages (1-2 and 7-10 for isoniazid, CCL4, Tetracyclines; and day 1-7, 12-14 for valproate) a true negative. Logistic regression with generalized linear model is performed using different combinations of PCs as predictors to maximize the probability of each datapoints belonging to its true classification for each drugs respectively, with formula:

$$Maximize\ \{logit(P(Y = true\_status)) = \beta 0 + \sum_{i+1}^{n} \beta i \cdot PCi\}$$

eventually we obtained as set of coefficients best separating the two classes pre-defined for each drug, with coefficients documented in table 1. By shifting threshold for class assignment, we get AUC-ROC of > 0.9 for isoniazid, tetracycline and valproate drugs respectively.

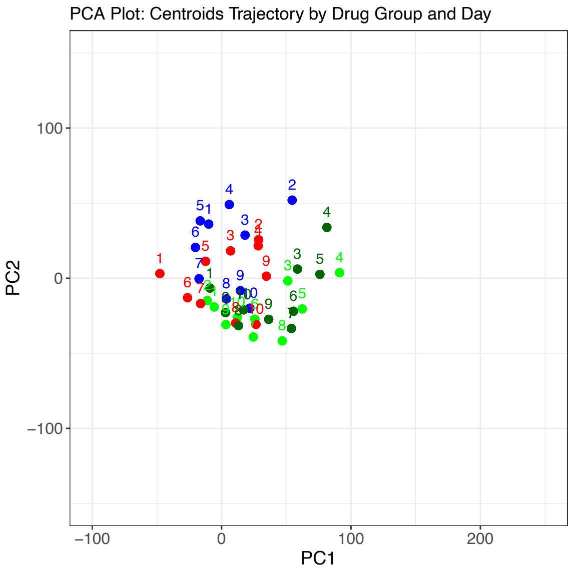


**Fig. S18** Centroid trajectories for subtoxic doses of all 4 drugs. Blue: Red: Light green: Dark Green:

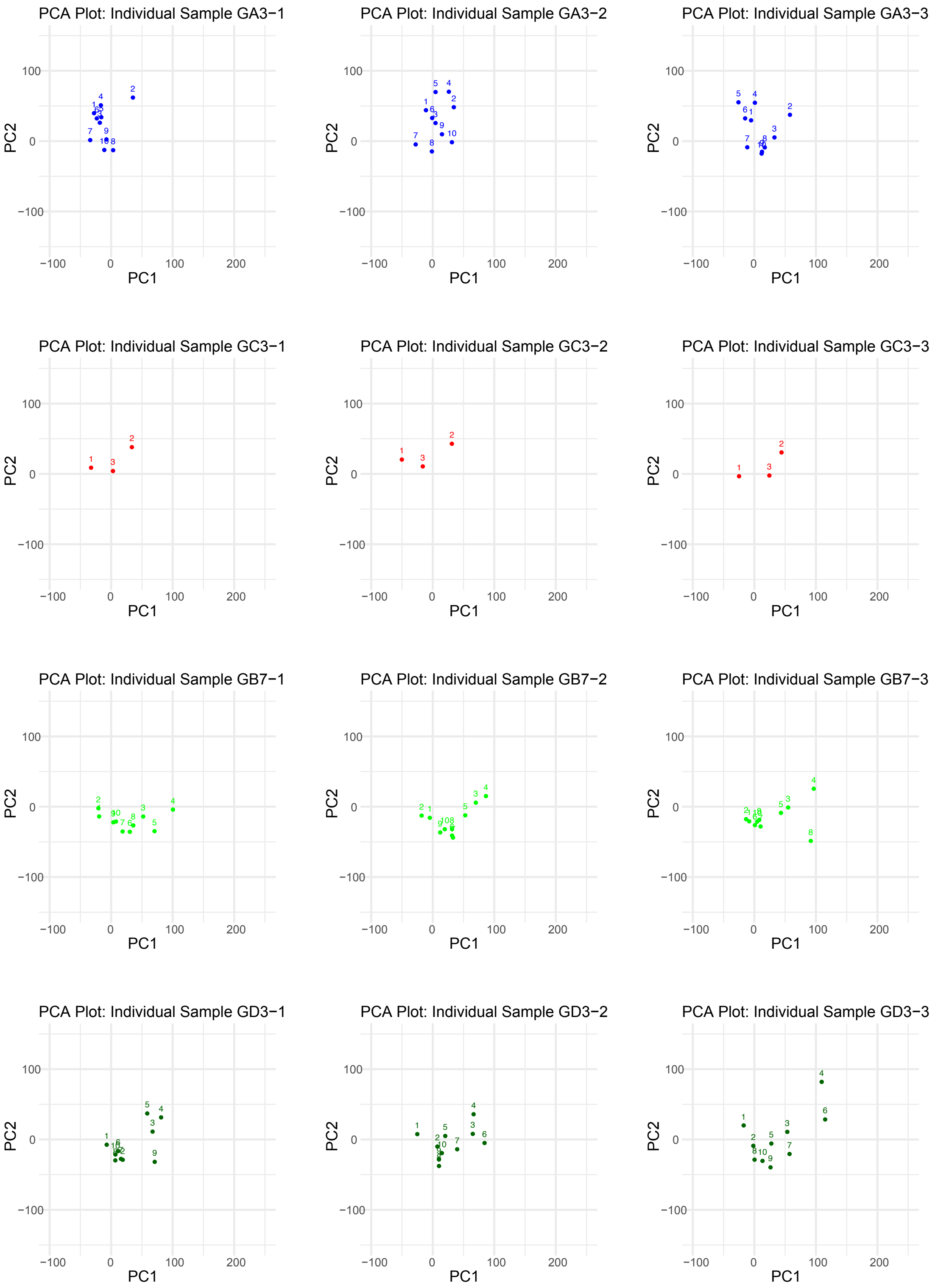


**Fig. S19** Sample individual trajectories for recovery to health

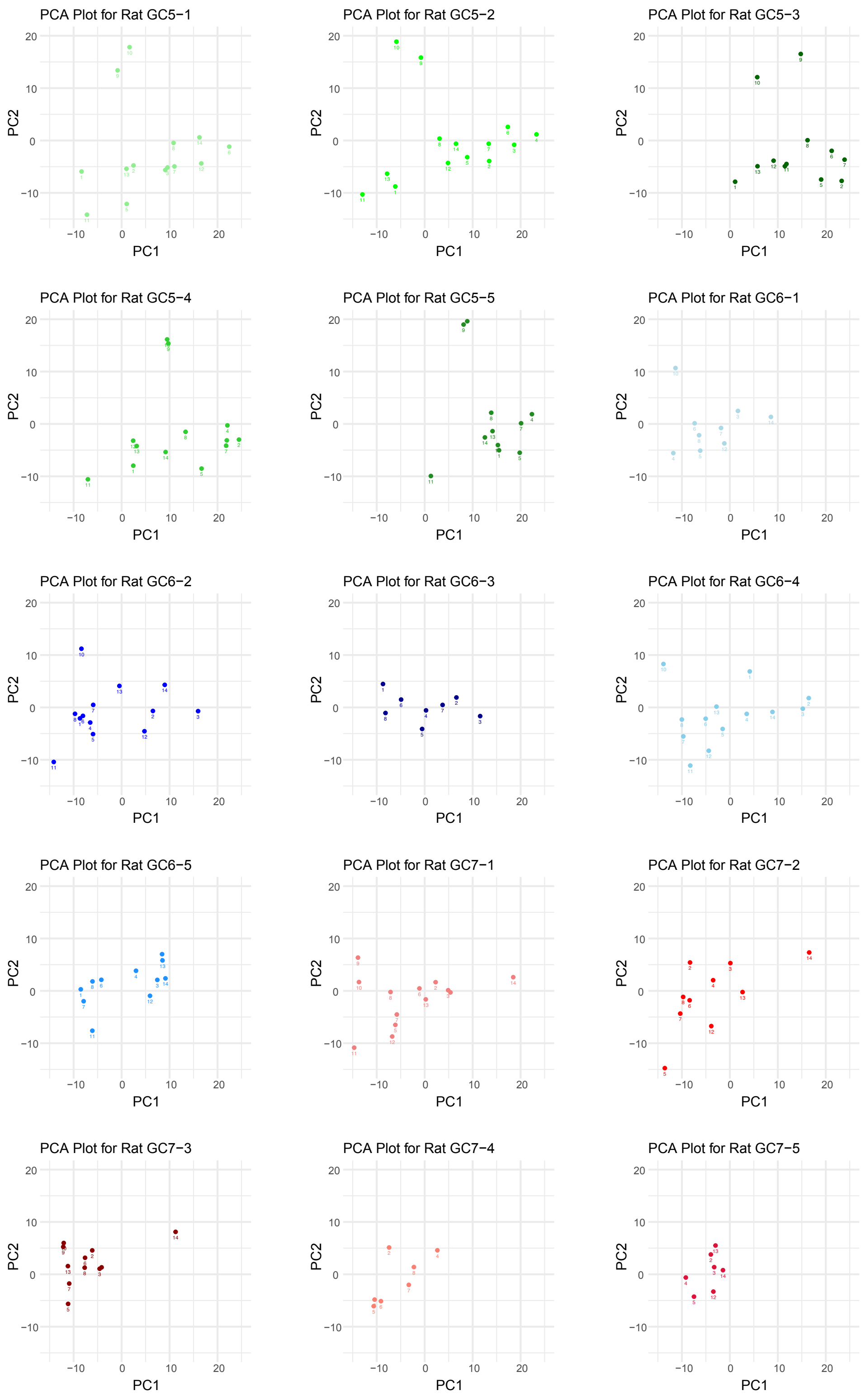


**Fig. S20** Trajectories in PC space for Fast Recovery, Slow Recovery

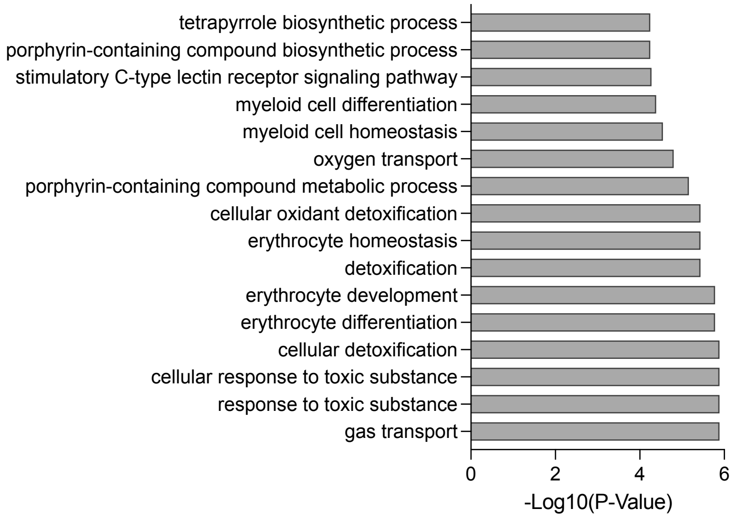


**Fig. S21** List of genes and pathway analysis for Fast Recovery genes.
